# JMJD1B-mediated FEN1 demethylation allows timely switching of Okazaki fragment maturation core enzymes to avoid mutagenic flap ligation by PARP1-LIG3

**DOI:** 10.1101/2025.10.06.680735

**Authors:** Yao Yan, Guojun Shi, Kejiao Li, Yi Lei, Yixing Wang, Yingying Wang, Songbai Liu, Anthony Fernandez, Tingting Zhou, Evan Wang, Mian Zhou, Yuan Hang, Zhuo Li, Zhigang Guo, Li Zheng, Binghui Shen

## Abstract

Efficient and faithful Okazaki fragment maturation (OFM) depends on the PCNA-coordinated actions of core enzymes Polδ, FEN1, and LIG1. We demonstrate that Polδ, FEN1, and LIG1 sequentially but not simultaneously bind to PCNA in mammalian cells. The association of FEN1 with PCNA, which lies at the center of this orderly program, is mediated by FEN1 R192 methylation, which is also crucial for preventing premature loading of LIG1. Conversely, FEN1 demethylation by the recently identified arginine demethylase JMJD1B promotes FEN1 dissociation from PCNA and LIG1 recruitment. Disruption of the sequential PCNA binding program in R192Q or *Jmjd1b^-/-^* cells results in unprocessed 5’ flaps that prevent OFM and induction of PARP1-dependent recruitment of LIG3, which has flap ligation activity to join incompletely processed OFs. This alternative OFM process supports cell survival but causes duplications and other DNA mutations. Our findings define fundamental and alternative DNA replication processes underlying mutagenesis and cell survival.

**Highlights:**

- The core enzymes involved in Okazaki fragment maturation (OFM), DNA polymerase δ (Polδ), flap endonuclease 1 (FEN1), and ligase 1 (LIG1), sequentially bind to the scaffold protein PCNA for their concerted actions
- FEN1 R192 methylation enables FEN1 binding to PCNA; JMJD1B-mediated FEN1 arginine demethylation provokes FEN1 exit from PCNA and LIG1 recruitment
- FEN1 R192Q mutation and JMJD1B deficiency oppositely alter dynamic PCNA interactions with FEN1 and LIG1 and result in distinct OFM DNA intermediate structures
- Activation of PARP1 at replication forks due to disruption of the sequential PCNA binding program induces LIG3-mediated alternative mutagenic 5’ flap ligation for survival.

## Introduction

Okazaki fragment maturation (OFM) is one of the most frequent DNA transactions. During lagging-strand DNA synthesis in mammalian cells, millions of Okazaki fragments (Sakofsky et al., 2014) are processed into an intact DNA strand in eukaryotic cells (Ma et al., 2022; Zheng and Shen, 2011). Unprocessed OFs, as single-strand breaks (SSBs), are potential sources of DNA double-strand breaks (DSBs), which are highly mutagenic and the most lethal type of DNA lesion. Thus, efficient and faithful OFM is of great importance to genome integrity and a critical determinant for cell survival. In recent decades, the key chemical reactions and corresponding core enzymes for OFM have been defined. DNA polymerase δ (Polδ) catalyzes DNA strand displacement synthesis, producing 5’ RNA-DNA primer flaps (Budd et al., 2006; Garg et al., 2004; Kahli et al., 2019; Nick McElhinny et al., 2008; Stith et al., 2008). Flap endonuclease 1 (FEN1) or the sequential actions of the helicase/nuclease DNA2 and FEN1 mediate nucleolytic degradation of 5’ flaps, generating ligatable DNA nicks (Ayyagari et al., 2003; Bae et al., 2001; Frank et al., 1998; Johnson et al., 1998; Liu et al., 2017; Tishkoff et al., 1997; Zheng et al., 2007a; Zheng et al., 2007b). DNA ligase 1 (LIG1) mediates ligation of the nick between two OFs to produce intact lagging strand DNA (Levin et al., 2000; Montecucco et al., 1998; Turchi et al., 1994; Waga et al., 1994). Due to the heavy workload of OFM in mammalian cells, it is important to optimize the actions of these three core enzymes for ordered, efficient, and faithful OFM.

The actions of Polδ, FEN1, and LIG1 are coordinated by proliferating cell nuclear antigen (PCNA), which forms a homo-trimeric structure encircling the DNA duplex, serving as a platform for recruitment of the core enzymes onto DNA replication forks (Chapados et al., 2004; Dore et al., 2006; Sporbert et al., 2005). Polδ, FEN1, and LIG1 share the same PCNA-binding motif, Qxx(L/I)xxF(F/Y) (Frank et al., 2001; Gary et al., 1997; Levin et al., 2000) for physical binding to a subunit of PCNA (Chapados et al., 2004; Dore et al., 2006; Refsland and Livingston, 2005; Sporbert et al., 2005; Subramanian et al., 2005). How PCNA coordinates the highly ordered actions of these core enzymes in the cell remains a mystery.

Two models were previously proposed to envisage how PCNA coordinates the actions of Polδ, FEN1, and LIG1 (Chapados et al., 2004; Dore et al., 2006; Refsland and Livingston, 2005; Sporbert et al., 2005; Subramanian et al., 2005). In one model, called the toolbelt model, Polδ, FEN1, and LIG1 simultaneously bind to each of the three subunits of the same DNA-bound PCNA trimer, which rotates along the DNA duplex (Blair et al., 2022; Dieckman et al., 2012; Lancey et al., 2020). The rotation allows Polδ, FEN1, and LIG1 to sequentially access intermediate DNA substrates for gap filling, flap cleavage, and nick ligation. However, LIG1 and FEN1 were shown to compete for PCNA (Subramanian et al., 2005), and resolution of the crystal structure of the LIG1-DNA complex indicated that LIG1 encircles the DNA substrate, excluding Polδ or FEN1 from interacting with PCNA (Pascal et al., 2004). These findings argue for mutually exclusive binding of PCNA with Polδ, FEN1, or LIG1 and suggests their interactions occur in a sequential order; this alternative model is called the sequential binding model, which is further by biochemical assays (Dovrat et al., 2014).

In the sequential binding model, competition for PCNA plays a crucial role in the orderly loading of the core enzymes onto PCNA. Changes in the PCNA-binding affinity of an enzyme, e.g., via post-translational modification, could enable such dynamic interactions with PCNA. Indeed, we previously showed that protein arginine methyltransferase 5 (PRMT5)-mediated arginine methylation and cyclin-dependent kinase 1/2 (Cdk1/2)-mediated phosphorylation of FEN1 mediate its interaction with PCNA during OFM (Guo et al., 2012; Guo et al., 2010; Xu et al., 2018). Methylation at the R192 residue of FEN1 promotes its interaction with PCNA and its recruitment to replication forks during OFM. In contrast, FEN1 S187 phosphorylation abolishes FEN1 interaction with PCNA, allowing LIG1 to bind PCNA. Interestingly, FEN1 S187 phosphorylation is dictated by FEN1 methylation status, as R192 FEN1 methylation changes FEN1 conformation and prevents Cdk1/2 from binding and phosphorylating FEN1 (Xu et al., 2018). Therefore, sequential FEN1 methylation, demethylation, and phosphorylation change FEN1 affinity for PCNA and preserve the FEN1-to-LIG1 transition at PCNA-encircled replication forks. However, the precise mechanism of PCNA-mediated coordination of all three core enzymes remains unresolved. One critical knowledge gap in this regulatory pathway is the identity of the arginine demethylase responsible for removal of R192 methylation from FEN1 as a key step for subsequent FEN1 phosphorylation and LIG1 recruitment.

Here, we report that PCNA primarily forms complexes with only one or two of the core enzymes at a time, forming a complex with Polδ, FEN1, and LIG1 only rarely. Thus, our studies suggest that the toolbelt model of simultaneous binding is not a major mechanism for PCNA to coordinate enzyme activities during OFM. Instead, Polδ, FEN1, or LIG1 sequentially interact with PCNA, forming two transient states, Polδ-PCNA-FEN1 and FEN1-PCNA-LIG1, during and after RNA-DNA 5’ flap processing, respectively. We further demonstrate that dynamic association of FEN1 with PCNA is crucial for maintaining the sequential PCNA binding program and that this dynamic association is mediated by FEN1 R192 methylation, catalyzed by PRMT5, and FEN1 R192 demethylation, which we now reveal is catalyzed by the histone arginine demethylase JMJD1B. FEN1 R192Q mutation reduces FEN1 stability and its association with PCNA and premature loading of LIG1. Conversely, the knockout of JMJD1B in MEFs results in FEN1 being trapped on PCNA at replication forks and subsequently delaying LIG1 from binding to PCNA for Okazaki fragment ligation. Both FEN1 R192Q and JMJD1B deletion results in OFM-related gaps and 5’ flaps and induces PARP1-dependent, error-prone alternative OF ligation by LIG3 for DNA replication.

## Results

### OFM core enzymes sequentially bind to the scaffold protein PCNA for their concerted reactions

To determine if PCNA binds Polδ, FEN1, and LIG1 simultaneously or sequentially during OFM, we first identified various PCNA-containing complexes in S phase cells. We treated HeLa cells with 1% formaldehyde to fix cells in S phase (when OFM occurs) and induce crosslinking, prepared nuclear extracts (NEs), and carried out immuno-blotting analyses without or with de-crosslinking by boiling the samples (Figure 1A). Using an anti-PCNA antibody, we identified three major bands corresponding to ∼36 kDa, ∼80 kDa, ∼160 kDa and a >200 kDa smear. De-crosslinking diminished all but the 36 kDa band, corresponding to the single PCNA subunit. It suggests that these bands greater than 36 kDa are protein complexes, not rather than modified forms of PCNA. Using anti-FEN1 antibody, in addition to a ∼43 kDa band, corresponding to the FEN1 monomer, we identified ∼80 kDa and ∼160 kDa bands and a >200 kDa smear as on the PCNA blot. Using an anti-LIG1 antibody, in addition to a ∼120 kDa band, indicative of the LIG1 monomer, we detected a ∼160 kDa band and >200 kDa smear as on the PCNA blot. Using anti-POLD1 (the catalytic and PCNA-binding subunit of Polδ) antibody, in addition to a ∼120 kDa band, indicative of the POLD1 monomer, we detected a unique ∼135 kDa band and >135 kDa smear. The molecular weight correspondence of protein bands in different immunoblots suggests that the ∼80 kDa PCNA band might contain the PCNA-FEN1 complex; the ∼160 kDa PCNA band might contain the PCNA-FEN1, the PCNA-LIG1, or the PCNA-POLD1 complex; and the > 200 kDa PCNA bands might contain the PCNA-FEN1-POLD1 or the PCNA-FEN1-LIG1 complex.

**Figure 1.**
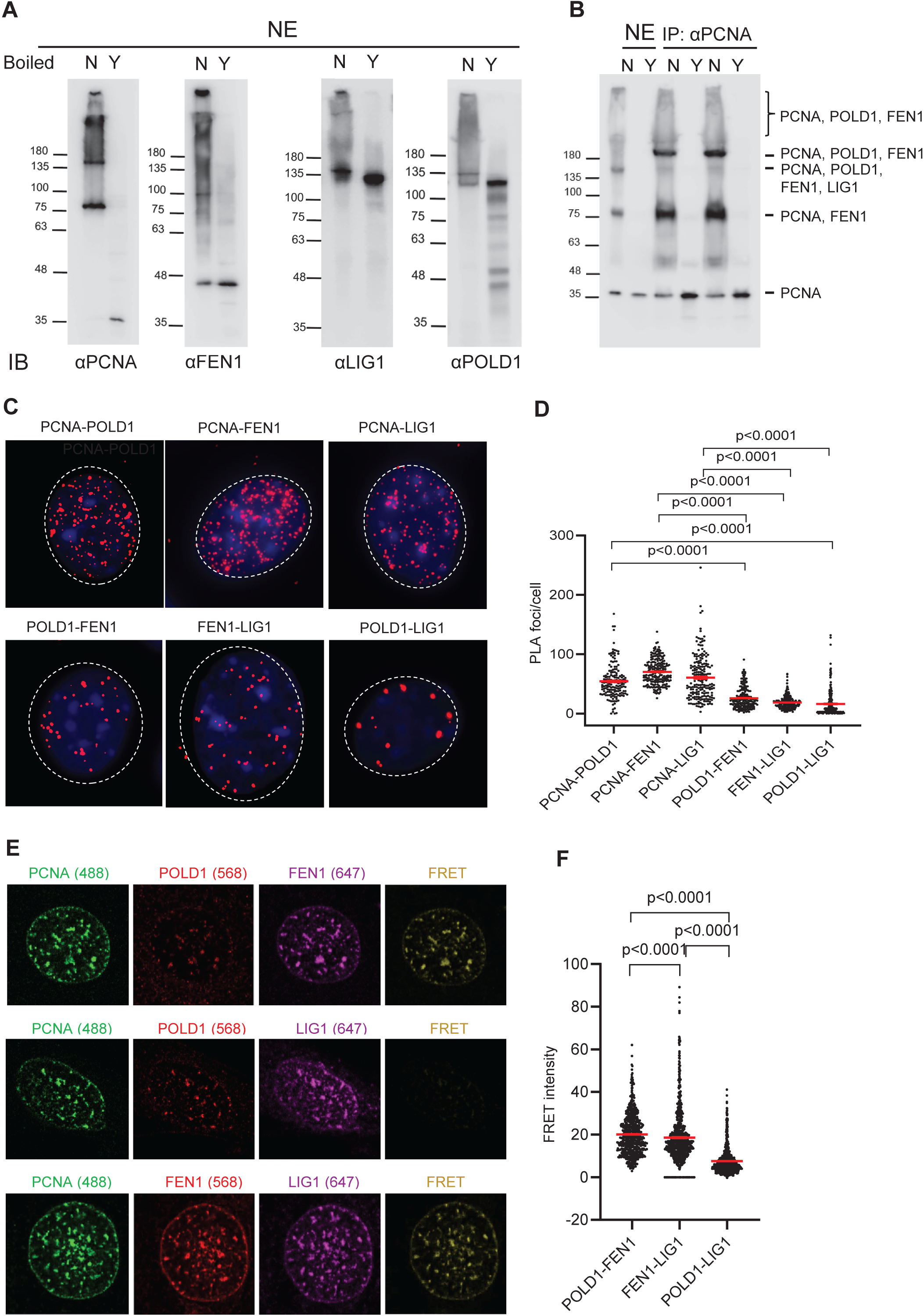
Dynamic PCNA complexes during OFM. **(A)** Non-boiled (N) and boiled (Y) nuclear extracts (NE) from HeLa cells after formaldehyde-crosslinking were analyzed by immuno-blot using indicated antibodies. **(B)** PCNA was IPed from HeLa NEs using anti-PCNA antibody and immuno-blot and mass spectrometry were used to identify the PCNA-bound proteins. The various PCNA-containing complexes are indicated. **(C)** Proximity ligation assay (PLA) was performed to detect *in situ* protein–protein interactions between PCNA-POLD1, PCNA-FEN1, PCNA-LIG1, POLD1-FEN1, FEN1-LIG1, and POLD1-LIG1 in WT MEFs. PLA signals (red puncta) were visualized using fluorescence microscopy (Zeiss Observer II). **(D)** Quantification of PLA signals per nucleus from panel C. Data represent mean ± SEM from ≥100 cells per condition. P value from Student t-test. **(E)** FRET analysis of protein–protein interactions between POLD1-FEN1, FEN1-LIG1, and POLD1-LIG1 in MEF WT cells. PCNA was used as a positional marker to indicate replication foci. Representative images for the control channel Alexa 488 (green; PCNA) and the FRET pair channels: Alexa 647 (purple), Alexa 568 (Blair et al., 2022), and FRET (yellow). **(F)** Quantification of the corrected FRET intensity from panel E. Data represent mean ± SEM from each foci per condition. P value from Student t-test.

To define the various PCNA-containing complexes, we first immunoprecipitated (IPed) PCNA from HeLa NEs (Figure 1B). In addition to ∼80 kDa and ∼160 kDa bands and a >200 kDa smear consistent with those in the anti-PCNA immuno-blot, a clear ∼200 band was enriched in the IPed sample. We then excised the bands and smears and performed mass spectrometry (not shown) to identify the following complexes: ∼80 kDa band: PCNA and FEN1; ∼160 kDa band: PCNA, POLD1, FEN1, and LIG1; ∼200 kDa band and >200 kDa smear: PCNA, POLD1, and FEN1. In PCNA complexes with molecular weight greater than ∼200 kDa, only FEN1 but not LIG1 was detected; this suggests the PCNA-POLD1-FEN1 complex might be more abundant and/or stable than the PCNA-FEN1-LIG1 complex.

To further define the various PCNA-containing complexes, we performed *in situ* experiments in mouse embryonic fibroblasts (MEFs). We previously used co-immunofluorescence (co-IF) (Cruz et al., 2018) staining to visualize FEN1 interaction with PCNA, as evidenced by co-localization. However, given that co-IF-detected PCNA foci are 0.5–1.0 μm in diameter and the various potential protein complexes are 1–100 nm in size, this method lacks the resolution needed to distinguish if Polδ, FEN1, and/or LIG1 bind to the same PCNA trimer or to two different PCNA trimers next to one another. To overcome this limitation, we employed Proximity Ligation Assay (PLA), which enables detection of endogenous protein– protein interactions *in situ* when targets are within close proximity (<40 nm) (Alam, 2018; Söderberg et al., 2008) to evaluate PCNA-containing complexes (Figure 1C and 1D). As expected, PCNA-POLD1, PCNA-FEN1, and PCNA-LIG1 protein pairs were the most common (∼40 PLA foci per MEF). In contrast, POLD1-FEN1 and FEN1-LIG1 (∼20 PLA foci per MEF) and POLD1-LIG1 (<5 PLA foci per MEF) protein pairs were significantly less common. This indicates that Polδ and LIG1 do not co-occur in the same PCNA complex, thus excluding the toolbelt model in which PCNA simultaneously binds Polδ, FEN1, and LIG1.

To further support this assertion, we performed Förster Resonance Energy Transfer (FRET) microscopy, which determines if two proteins physically interact with one another by detecting the transfer of energy from a donor fluorophore to an acceptor fluorophore that is typically <10 nm away from the donor fluorophore (Sekar and Periasamy, 2003). We evaluated FRET signals to determine whether Polδ and FEN1, Polδ and LIG1, and/or FEN1 and LIG1 co-occur in the same PCNA trimer (Figure 1E and 1F). We detected PCNA foci-colocalized FRET signal between POLD1 and FEN1 (20.1±0.4) or between FEN1 and LIG1 (18.6±0.5) However, we detected significantly less FRET signal between POLD1 and LIG1 (7.5±0.2). This is consistent with results of both the PCNA IP and PLA experiments.

Collectively, these data support a sequential binding model for PCNA during OFM. PCNA first forms a complex with Polδ for primer extension, then recruits FEN1, leading to a PCNA-Polδ-FEN1 complex for strand displacement DNA synthesis and 5’ flap cleavage, as suggested previously in a biochemical assay (Matsumoto et al., 2020). Polδ then dissociates, leaving a PCNA-FEN1 complex, which likely mediates the final round of 5’ flap cleavage. PCNA then recruits LIG1, producing a transient PCNA-FEN1-LIG1 complex; subsequent FEN1 exit leads to a PCNA-LIG1 complex for OF ligation. The transient trimeric complexes (PCNA-Polδ-FEN1 and PCNA-FEN1-LIG1) exist for only a short time as the proteins are switched. The limited detection of a Polδ-LIG1 complex suggests that a fully assembled PCNA-Polδ-FEN1-LIG1 complex may exist only as a transient and unstable intermediate during OFM.

### JMJD1B mediates FEN1 arginine demethylation during OFM

An important question is how Polδ, FEN1, and LIG1 are dynamically loaded onto and unloaded from the PCNA platform. FEN1 lies at the center of the sequential PCNA binding program. We therefore assert that dynamic interaction between FEN1 and PCNA is crucial not only for processing 5’ flaps of RNA-DNA primers but also important for timely unloading of Polδ from PCNA or loading of LIG1 onto PCNA. Furthermore, our previous studies suggest that dynamic interaction of FEN1 with PCNA is controlled by the interplay between FEN1 R192 methylation and S187 phosphorylation (Guo et al., 2012; Guo et al., 2010). We and others previously showed that FEN1 S187 phosphorylation, which occurs in late S phase, abolishes FEN1 interaction with PCNA and triggers its unloading from replication sites and eventual degradation. It is important to note that FEN1 phosphorylation is suppressed by FEN1 R192 methylation, which induces a conformation that blocks Cdk1 from binding and phosphorylating FEN1. This suggests that in early S phase, FEN1 R192 is methylated to ensure PCNA binding, but in late S phase R192-methylated FEN1 is demethylated to allow FEN1 phosphorylation and dissociation from PCNA.

To test this hypothesis, we synchronized HeLa cells at the G1/S boundary, released the cells into S phase, then IPed FEN1 and performed immuno-blotting. We observed that relative to total FEN1, methylated FEN1 increased considerably by 4 h post-G1/S boundary but decreased by 8 h post-G1/S boundary (Figure 2A). This observation is consistent with the hypothesis that FEN1 is methylated in early S phase and demethylated in late S phase. We previously showed that FEN1 arginine methylation is catalyzed by PRMT5 (Guo et al., 2010) However, the arginine demethylase for FEN1 is unknown. To identify the FEN1 demethylase, we IPed FEN1 protein complexes from NEs of HeLa cells, then performed LC-MS/MS analysis to identify potential FEN1-interacting proteins. In addition to known FEN1-interacting proteins, we found that JMJD1B, a Jumonji C domain-containing protein, interacted with FEN1 (Supplementary Table S1). We used immuno-blot to confirm that JMJD1B co-IPed with FEN1 in NEs from S phase HeLa cells (Figure 2B). Of note, the amount of JMJD1B that co-IPed with FEN1 greatly increased at 4 h post-G1/S boundary (Figure 2B) but decreased at 8 h post-G1/S boundary, concurrent with the reduction in methylated FEN1. This suggests that JMJD1B interacts with methylated FEN1. To confirm this, we used a co-pull-down assay to demonstrate that purified recombinant JMJD1B and FEN1 proteins physically interacted with one another (Figure 2C). In addition, we found that deletion of the FEN1 C-terminal region important for FEN1 interactions with many proteins (Guo et al., 2009) also abolished FEN1 interaction with JMJD1B (Supplementary Figure S1A).

**Figure 2.**
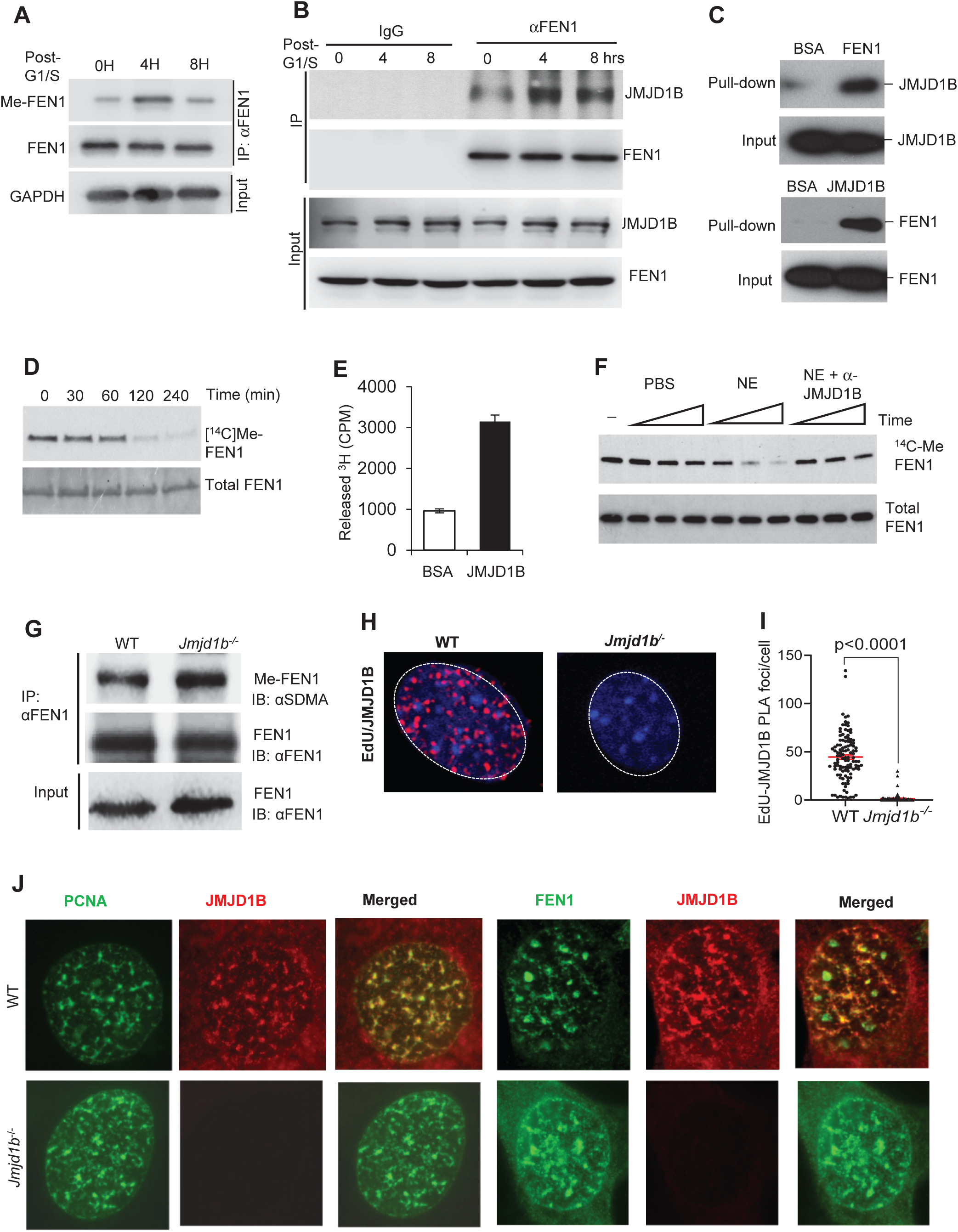
JMJD1B interacts with and demethylates FEN1 in S phase. **(A)** Methylated FEN1 (MeFEN1) protein in HeLa cells. HeLa cells were synchronized at the G1/S boundary (using mimosine) and released into S phase for 0, 4 or 8 h. FEN1 was pulled down from NEs, meFEN1 and total FEN1 were detected using immuno-blotting. **(B)** JMJD1B that was co-IPed with FEN1 from S phase HeLa NEs was detected using immuno-blotting (IB). **(C)** Co-pull-down assays using purified FEN1 and JMJD1B show physical interaction between FEN1 and JMJD1B. **(D)** ^14^C-labeled methylated FEN1 was incubated with purified recombinant JMJD1B (active motif) at 37°C for 60, 120, or 240 min. The samples were subjected to autoradiography analysis of ^14^C-meFEN1. **(E)** ^3^H-labeled methylated FEN1 was incubated with purified recombinant JMJD1B (active motif) at 37°C for 2 h. Free ^3^H released from ^3^H-meFEN1 was measured using a scintillation counter. Values are means ± s.d. of three independent assays. **(F)** ^14^C-labeled methylated FEN1 was incubated with HeLa NEs with or without JMJD1B antibody depletion at 37°C for 1, 2, or 4 h. Methylated FEN1 was detected using autoradiography and total FEN1 level was detected using immuno-blotting. **(G)** Total SDMA-modified proteins isolated from WT or *Jmjd1b*^-/-^ MEFs were IPed with a rabbit anti-synmetric dimethylation (SDMA) antibody (Millipore). meFEN1 that was pulled down by the anti-SDMA antibody was analyzed by immuno-blot using a mouse monoclonal anti-FEN1 antibody (Genetex). **(H)** PLA was performed to detect JMJD1B in newly synthesized DNA in WT MEFs and *Jmjd1b*^-/-^ MEFs. **(I)** Quantification of PLA signals per cell from panel H. Data represent means ± SEMs from ≥100 cells per condition. P value from Student t-test. **(J)** Immunostaining assay was applied to detect the colocalization between PCNA and JMJD1B, FEN1 and JMJD1B in WT MEFs and *Jmjd1b*^-/-^ MEFs.

To determine if JMJD1B catalyzes demethylation of arginine methylated FEN1 (meFEN1), we incubated purified recombinant JMJD1B with ^14^C-labeled or ^3^H-labeled meFEN1. We observed a decrease in ^14^C-labeled meFEN1 (Figure 2D) and a concomitant increase in production of free ^3^H-methyl group (Figure 2E). This demonstrates that meFEN1 is a substrate of JMJD1B arginine demethylase. To determine if JMJD1B is the primary arginine demethylase for FEN1 in cells, we prepared NEs from HeLa cells, depleted JMJD1B using an antibody, and incubated NEs with ^14^C-labeled meFEN1. Depletion of JMJD1B from HeLa NEs notably reduced demethylation of ^14^C-labeled meFEN1 (Figure 2F), suggesting JMJD1B is the primary FEN1 demethylase in HeLa cells. To further confirm that JMJD1B demethylates FEN1 *in vivo*, we measured meFEN1 in MEFs from wild-type (WT) and Jmjd1b knockout (*Jmjd1b*^-/-^) mice, a model we previously established (Li et al., 2018). We found that relative to WT MEFs, *Jmjd1b*^-/-^ MEFs exhibited considerably increased level of meFEN1 (Figure 2G). Together, these data demonstrate that JMJD1B acts as a FEN1 arginine demethylase. Given that JMJD1B demethylates meFEN1 during OFM, we would expect to detect JMJD1B at replication forks. To confirm that JMJD1B associates with replication forks, we labeled the cells with the nucleoside analog EdU, which is used as a readout of replication forks (Abe et al., 2018) and performed PLA for EdU and JMJD1B. We detected on average 44.7±2.3 JMJD1B-EDU PLA foci per nucleus in WT MEF cells but almost none (1.5±0.5/nucleus) in the negative control (*Jmjd1b*^-/-^ MEFs) (Figure 3H and 3I). In addition, we carried out co-IF staining of JMJD1B and PCNA, as a molecular marker for replication forks, and JMJD1B and FEN1 in MEF and HeLa cells. We showed that JMJD1B formed foci that co-localized with PCNA foci and FEN1 foci (Figure 2J, Supplementary Figure S1B).

**Figure 3.**
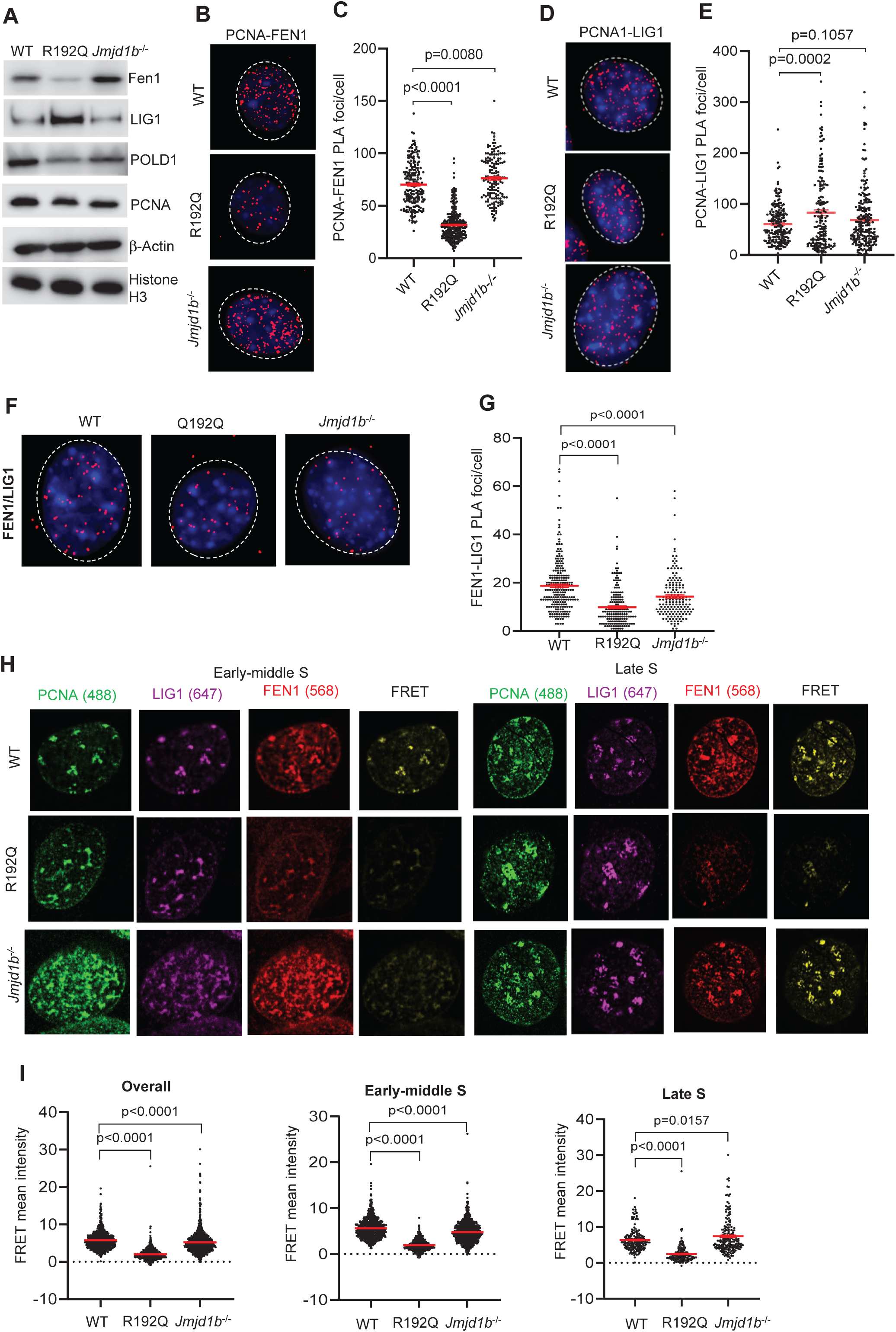
Disruption of FEN1 arginine methylation/demethylation impairs the dynamic interaction of FEN1 with PCNA and the sequential PCNA binding program. **(A)** Immuno-blot of whole cell lysates for FEN1, LIG1, POLD1, PCNA, β-Actin and Histone H3 in WT, FEN1 R192Q, and *Jmjd1b*^-/-^ MEFs. **(B-G)** PLA was performed to detect interaction between PCNA-FEN1 (B, C), PCNA-LIG1 (D, E) and FEN1-LIG1 (F, G) in WT, FEN1 R192Q, and *Jmjd1b*^-/-^ MEFs. PLA signals (red puncta) were visualized using fluorescence microscopy (Zeiss Observer II). Panel B, D, or F shows the representative PLA images and panel C, E, or G shows quantification of PLA signals per cell from panel B, D, and F. Data represent mean ± SEM from ≥100 cells per condition. P value from Student t-test. **(H, I)** FEN1-LIG1 FRET was performed to assess the overall level of the FEN1-LIG1 complex at various stages of S phase in in WT, R192Q, or *Jmjd1b^-/-^* MEFs. (H) Representative images for the Alexa 488 (PCNA; green), Alexa 647 (LIG1; purple), Alexa 568 (FEN1; red), or FRET (yellow) channel. The different stages of S phase were defined based on the pattern of PCNA foci. Early: relatively uniform distribution of small PCNA foci; middle: PCNA foci with increased numbers and intensity present in the nuclei and nuclear membranes; late: large clusters of PCNA foci in nuclei. (I) Quantification of corrected FRET intensity from panel H. Data represent means ± SEMs. P value from Student t-test.

### FEN1 methylation and demethylation control OFM core enzymes sequential interactions with PCNA

We next investigated the biological importance of dynamic FEN1 R192 methylation and demethylation on FEN1 dynamics at the PCNA platform. We previously reported a human FEN1 single nucleotide polymorphism R192Q (rs4989586) that eliminates the methylation site (Guo et al., 2010). To assess the impact of FEN1 R192Q mutation or *Jmjd1b* deficiency on the dynamic interactions of FEN1 with PCNA during DNA replication, we performed immuno-blot. We observed that FEN1 expression was greatly reduced in R192Q MEFs, compared to WT or *Jmjd1b^-/-^* MEFs (Figure 3A). This observation is consistent with previous findings that defects in FEN1 R192 methylation promote FEN1 phosphorylation and degradation (Guo et al., 2012). We further assessed the interaction of FEN1 and PCNA using FEN1-PCNA PLA, co-IF staining, and FRET in WT, R192Q, and *Jmjd1b^-/-^* MEFs. Compared to WT MEFs (70.3±1.4 foci/nucleus), the number of FEN1-PCNA PLA foci was significantly lower in R192Q MEFs (∼31.7±0.8 foci/nucleus) but significantly greater in *Jmjd1b^-/-^*MEFs (∼76.2±1.7 foci/nucleus) (Figure 3B and 3C). Consistent with this, the intensity of both PCNA-colocalized FEN1 foci (Supplementary Figure S2A and S2B) and FEN1-PCNA FRET (Supplementary Figure S3A and S3B) were significantly lower in R192Q MEFs but significantly greater in *Jmjd1b^-/-^* MEFs than in WT MEFs.

These findings suggest that FEN1 R192 methylation is crucial for FEN1 stability and interaction with PCNA, whereas demethylation of meFEN1 by JMJD1B is important for FEN1 dissociation from PCNA. In addition, we noted that in early-to-middle S phase, PCNA foci were uniformly co-localized with FEN1 foci in both WT and *Jmjd1b*^-/-^ MEFs (Supplementary Figure S2C and S2D). However, in late S phase, in which PCNA foci form clusters as previously defined (Essers et al., 2005; Refsland and Livingston, 2005; Zerjatke et al., 2017) co-localization of PCNA foci with FEN1 foci, and the corresponding intensity of FEN1 foci, were greatly reduced in WT MEFs compared to *Jmjd1b*^-/-^ MEFs (Supplementary Figure S2C, S2D). This finding further supports the importance of JMJD1B-mediated FEN1 arginine demethylation for FEN1 dissociation from PCNA.

FEN1 lies at the center of the sequential PCNA binding program; indeed, we observed that FEN1 co-localization with PCNA was inversely associated with LIG1 co-localization with PCNA (Supplementary Figure S4). We therefore hypothesized that disruption of FEN1 R192 methylation and demethylation, which alters the dynamic interaction of FEN1 with PCNA, affects LIG1 binding to PCNA for OF ligation. To test this, we used PLA to examine PCNA interaction with LIG1 in WT, R192Q, and *Jmjd1b^-/-^* MEFs. We found that the number of PCNA-LIG1 PLA foci was significantly greater in R192Q MEFs (82.9±5.7 foci/nucleus) than in WT MEFs (60.6±2.7foci/nucleus) (Figure 3D, 3E). This suggests that retardation of FEN1 R192Q recruitment leads to premature loading of LIG1 onto PCNA. We also observed significantly fewer FEN1-LIG1 PLA foci in R192Q MEFs (9.8±0.6 foci/nucleus) than in WT MEFs (∼18.8±0.7 foci/nucleus) (Figure 3F, 3G). Using FRET microscopy to visualize co-localization of FEN1-LIG1 with PCNA foci in early-middle and late S phase, we observed significantly lower FEN1-LIG1 FRET levels at PCNA foci in R192Q MEFs (2.0±0.1) than in WT MEFs (5.8±0.1) throughout S phase (Figure 3H, 3I).In contrast to what observed in R192Q cells, we did not detect significantly more PCNA-LIG1 PLA foci per cell in *Jmjd1b-/-* cells than in the WT cells (Figure 3D, 3E). Instead, we observed considerably less PCNA-colocalized LIG1 in early middle S phase in *Jmjd1b-/-* MEFs than in WT MEFs (Supplementary Figure S5A and S5B). This suggests that persistent association of FEN1 with PCNA in *Jmjd1b*^-/-^ MEFs at least partially impairs LIG1 recruitment. Intriguingly, as *Jmjd1b*^-/-^ MEFs progressed to the late stage of S phase, we observed PCNA-colocalized LIG1 foci similar to those in WT MEFs (Supplementary Figure S5A and S5B), even though FEN1 remained associated with PCNA. Consistent with this, we observed a significant decrease in PCNA-co-localized FEN1-LIG1 FRET signals in *Jmjd1b*^-/-^ MEFs (4.8±0.1) compared to WT MEFs (5.6±0.1) in early middle S phase, reflecting defective LIG1 recruitment to PCNA (Figure 3H, 3I). However, in late S phase, we observed a greater FEN1-LIG1 FRET signal at PCNA foci in *Jmjd1b*^-/-^ MEFs (7.4±0.4) than in WT MEFs (6.3±0.2) (Figure 3H, 3I), indicating increased levels of PCNA-FEN1-LIG1 in *Jmjd1b*^-/-^ vs. WT MEFs.

### Disruption of FEN1 methylation dynamics results in replicative DNA intermediates, DSBs, and chromosome segregation abnormality

To define the impact of FEN1 R192Q mutation or JMJD1B deficiency on OFM efficiency, we used a BrdU alkaline Comet assay and measured replicative ssDNA breaks in WT, R192Q, and *Jmjd1b^-/-^* MEFs. On average, BrdU-labeled comet DNA tail moment (tail length in nm x % tail DNA) was increased in R192Q (218.5±11.5) and *Jmjd1b^-/-^* (221.3±9.2) compared to WT (83.6±6.2) MEFs (Figure 4A, 4B), indicative of accumulation of replicative ssDNA breaks. To determine which specific alteration occurred on replication forks in R192Q and *Jmjd1b^-/-^* vs. WT MEFs, we used transmission electron microscopy (TEM) to analyze DNA replication fork intermediate structures *in vivo*. We enriched and purified replication forks, then performed benzalkonium chloride (BAC)-mediated spreading and direct visualization using TEM. Consistent with previous observations on replication forks from yeast cells, in MEFs we observed Y-shaped structures of replication forks, which contained a double-stranded parental strand and two double-stranded daughter strands of equal length (Figure 4C-4E). The presence of intermediate structures (ssDNA flaps and gaps) was significantly increased in replication forks in R192Q MEFs (57.4% ssDNA flaps, 29.5% ssDNA gaps) and *Jmjd1b^-/-^* MEFs (35.8% ssDNA flaps, 44.8% ssDNA gaps) compared to those in WT MEFs (12.1% ssDNA flaps, 5.4% ssDNA gaps) (Figure 4F, 4G). In addition, two or three ssDNA gaps on a single daughter strand occurred more frequently in *Jmjd1b^-/-^* MEFs (∼20%) than in R192Q (∼1%) or WT (0%) MEFs (Figure 4G). There was no significant difference in the mean length of unprocessed ssDNA flaps in R192Q (57 ± 35 nt) or *Jmjd1b^-/-^* (50 ± 19 nt) compared to WT (42 ± 21 nt) MEFs (Figure 4H). In contrast, the mean length of ssDNA gaps was significantly increased in R192Q (108 ± 37 nt) and *Jmjd1b^-/-^* (142 ± 89 nt) compared to WT (54 ± 28 nt) MEFs (Figure 4I). These findings indicate that defects in FEN1 R192 methylation due to R192Q mutation or in demethylation due to JMJD1B deficiency caused accumulation of ssDNA flaps and gaps at replication forks. Of note, FEN1 R192Q mutation, which causes ssDNA flap processing, resulted in greater ssDNA flap accumulation, whereas JMJD1B deficiency, which causes persistent FEN1 association with PCNA and delays LIG1 recruitment, resulted in greater ssDNA gap accumulation. Our observation that defective LIG1 recruitment causes 5’ flap accumulation in *Jmjd1b^-/-^* MEFs is consistent with a previous report showing that conditional depletion of DNA ligase Cdc9 in *Saccharomyces cerevisiae* leads to an accumulation of flap structures on the lagging strands of replication forks (Shi et al., 2024).

**Figure 4.**
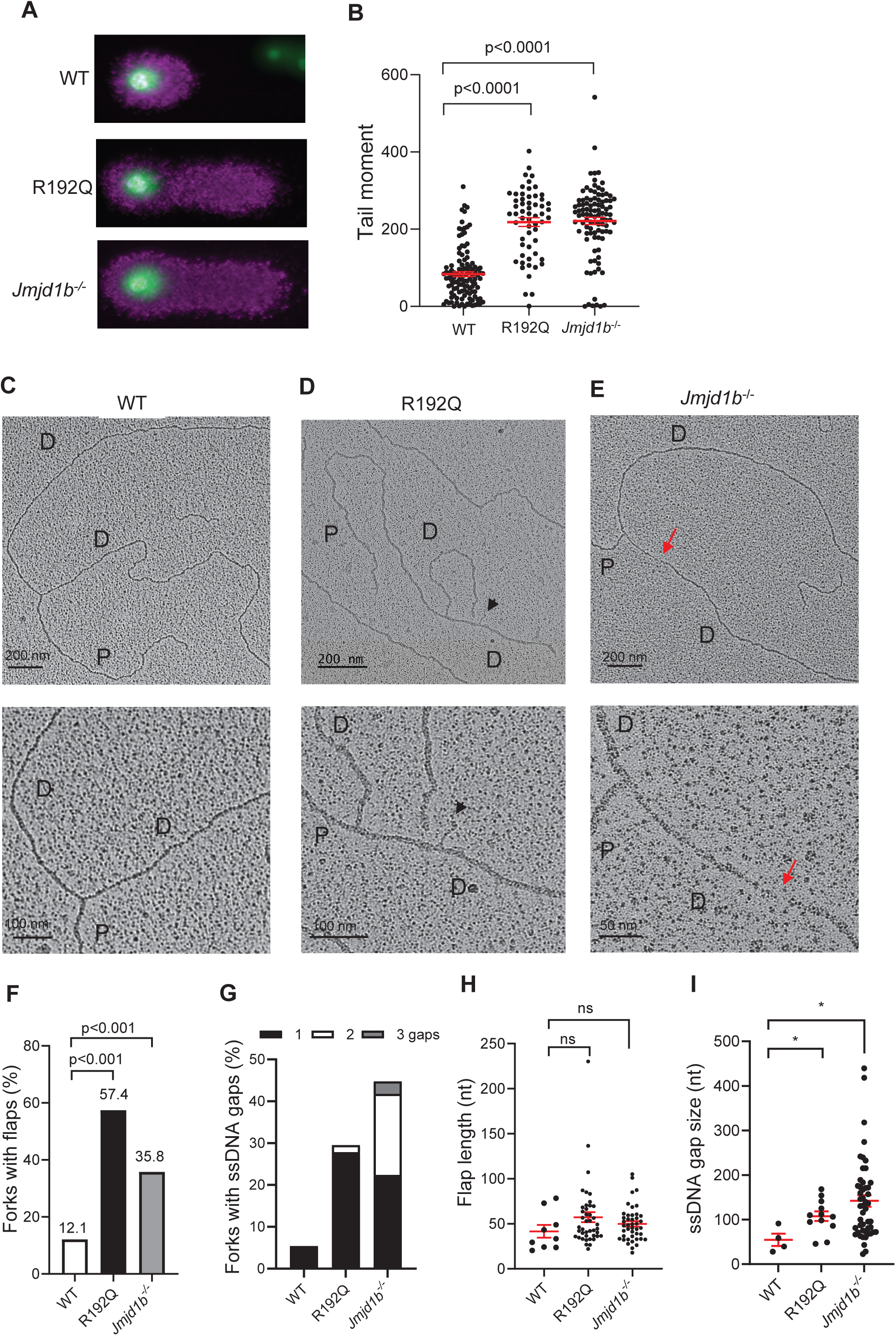
FEN1 R192Q mutation or JMJD1B deficiency impairs OFM and causes accumulation of 5’ flaps and gaps at replication forks. **(A-B)** Representative fluorescence microscopy images (A) and quantification of tail moment (B) from the Alkaline-BrdU comet assay performed on WT, FEN1 R192Q, and *Jmjd1b^-/-^* MEFs. Cells were pulse-labeled with 20 μM BrdU for 20 min prior to experiment. For each sample, at least 50 comets were quantified. Mean ± SEM. are indicated. P value from Student t-test. **(C-E)** Representative TEM images of replication forks in WT, R192Q, and *Jmjd1b^-/-^*MEFs. D, daughter strand; P, parental strand. Scale bars, 200 nm [500 base pairs (bp)]. Black arrowheads point to ssDNA flaps; red arrows point to ssDNA gaps. The enlarged views are shown below the image, with scale bars as indicated. **(F-G)** Percentage of replication forks with flaps (F) or with 1, 2, or 3 ssDNA gaps (G) in WT (n=74 total observed forks), R192Q (n=61), and *Jmjd1b^-/-^* (n=67) MEFs. **(H)** Average length of ssDNA flaps in WT (n=9 total observed flaps), R192Q (n=42), and *Jmjd1b^-/-^*(n=43) MEFs from two independent experiments. Bars represent means ± SEMs. P value from Student t-test. **(I)** Average length of ssDNA gaps in WT (n=4 total observed gaps), R192Q (n=12), and *Jmjd1b^-/-^* (n=49) MEFs from two independent experiments. Bars represent means ± SEMs. P value from Student t-test.

To determine if accumulated replicative ssDNA breaks including DNA nicks and gaps with or without flaps are converted into DSBs in R192Q or *Jmjd1b*^-/-^ MEFs, we performed neutral comet assays, which detect genomic DSBs at the single-cell level. We found that tail moment was significantly greater in R192Q (9.3±0.6) and *Jmjd1b*^-/-^ (15.0±0.8) MEFs than in WT MEFs (2.0±0.2) (Figure 5A and 5B). Consistent with this, we observed that R192Q (13.9±1.2 foci/nucleus) and *Jmjd1b*^-/-^ (15.06±1.5/nucleus) MEFs exhibited significantly more γH2AX foci, a marker for histone response to DSBs (Refsland and Livingston, 2005), than did WT MEFs (8.2±0.7 foci/nucleus) (Figure 5C and 5D). Similarly, R192Q (5.8±0.8 foci/nucleus) and *Jmjd1b^-/-^*(12.0±0.9 foci/nucleus) MEFs exhibited significantly more RAD51 foci, a marker for activation of homology-directed repair of DSBs in S or G2 phase cells (Cruz et al., 2018), than did WT MEFs (3.3±0.3 foci/nucleus) (Figure 5C and 5E). We also observed significantly more frequent chromosome segregation errors (e.g., bridging, lagging) in anaphase in R192Q and *Jmjd1b*^-/-^ mutant MEFs than in WT MEFs (Figure 5F and 5G). Together, these results indicate that disruption of FEN1 methylation and demethylation dynamics changes the sequential PCNA binding program and results in replicative ssDNA gaps, flaps, and DSBs and frequent chromosome segregation errors.

**Figure 5.**
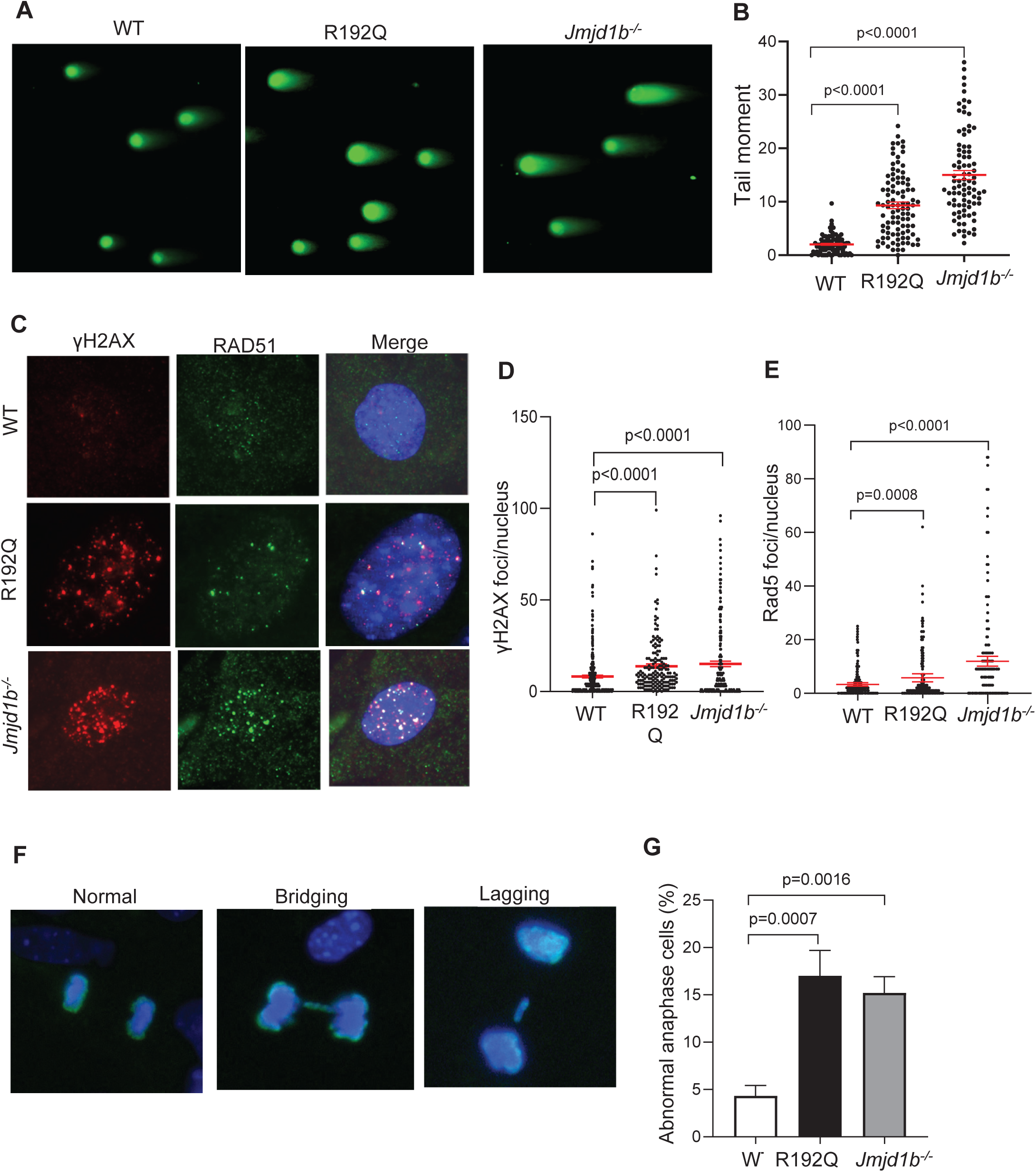
FEN1 R192Q mutation or JMJD1B deficiency induces accumulation of DSBs and DNA mutations and aberrant chromosome segregation. **(A-B)** Representative microscopy images (A) and quantification of tail moments (B) from neutral comet assay performed on WT, FEN1 R192Q, and *Jmjd1b^-/-^* MEFs. At least 50 comets were quantified per sample. Mean ± SEM. are indicated. P value from Student t-test. **(C-E)** γH2AX and RAD51 foci in WT, FEN1 R192Q, and *Jmjd1b*^-/-^ MEFs. Representative Co-IF staining images of γH2AX (Blair et al., 2022) and RAD51 (green) (C) in WT, FEN1 R192Q, and *Jmjd1b*^-/-^ MEFs. Quantification of γH2AX (D) and RAD51 (E) foci in WT, FEN1 R192Q, and *Jmjd1b^-/-^* MEFs using the Image J program. Mean ± S.E.M. are indicated. P value from Student t-test. **(F-G)** Representative microscopy images (F) of indicated chromosome segregation errors and quantification of abnormal anaphase cells (G) in WT, FEN1 R192Q, and *Jmjd1b^-/-^* MEFs stained with phosphor-histone H3 (Ser10) (green) and DAPI (blue). Means ± SEMs are indicated. P value from Student t-test.

### Defective canonic OFM induces PARP1-LIG3-dependent mutagenic OF ligation for cell survival

As shown above, defects in FEN1 methylation or demethylation impaired flap processing and OF ligation and resulted in un-ligated OFs in the genome. To determine if R192Q FEN1 mutation and *Jmjd1b^-/-^* MEF cells induce alternative pathways for OFM to compensate the defects, we have examined the expression levels of LIG1, DNA ligase 3 (LIG3) and PARP1 genes. We showed that LIG1 was overexpressed (Figure 3A) and pre-maturely loaded onto PCNA in R192Q mutant, but not WT, MEF cells (Figure 3B, 3D). A previous study showed that defects in OFM due to FEN1 deficiency or inhibition results in PARP1 chromatin accumulation and recruitment to replication forks (Hanzlikova et al., 2018). Indeed, we observed more chromatin-associated PARP1 in R192Q MEFs than in the WT (Figure 6A) and detected significantly more PARP1-EdU PLA foci in R192Q MEFs (10.7±1.8 foci/nucleus) than in WT (4.7±1.2) (Figure 6B, 6C). Meanwhile, we found that the level of chromatin-associated PARP1 in *Jmjd1b^-/-^* MEFs was considerably higher than in the WT (Figure 6A). PARP1-EdU PLA foci in *Jmjd1b^-/-^*MEFs (9.8±1.5 foci/nucleus) were significantly more than that in the WT (Figure 6B, 6C). Given that PARP1 activation at SSBs promotes recruitment of LIG3 is thought to back up the function of LIG1 (Arakawa et al., 2012; Refsland and Livingston, 2005), we used immuno-blot and EdU-LIG3 PLA to assess if SSBs in R192Q or *Jmjd1b^-/-^* MEFs increases chromatin-associated and/or induces LIG3 binding to replication forks. We observed elevated chromatin-associated LIG3 in both R192Q and *Jmjd1b^-/-^* compared to WT MEFs (Figure 6A). Furthermore, EdU-LIG3 foci were significantly increased in R192Q (18.0±2.5 PLA foci/nucleus) and *Jmjd1b^-/-^*(21.8±2.6 PLA foci/nucleus) MEFs compared to WT MEFs (3.9±0.9 PLA foci/nucleus) (Figure 6D, 6E). To determine whether PARP1 activity is important for recruiting LIG3 to replication forks, we inhibited PARP1 activity using Olaparib (1 or 10 μM, 16 h), which significantly reduced the number of EdU-LIG3 foci in R192Q or *Jmjd1b^-/-^* MEFs, similar to that in the untreated WT control (Figure 6D, 6E). These findings suggest that defects in FEN1 binding to PCNA in R192Q MEFs or defects in the FEN1-to-LIG1 transition in *Jmjd1b^-/-^* MEFs result in PARP1-dependent recruitment of LIG3 to replication forks to mediate alternative OF ligation. Consistent with this, we observed that Olaparib (1 or 10 μM, 16 h) significantly increased the level of replicative SSBs, γH2AX foci, and RAD51 foci in WT, R192Q, and *Jmjd1b^-/-^*MEFs (Figure 6F, 6G, Supplementary Figure S6).

**Figure 6.**
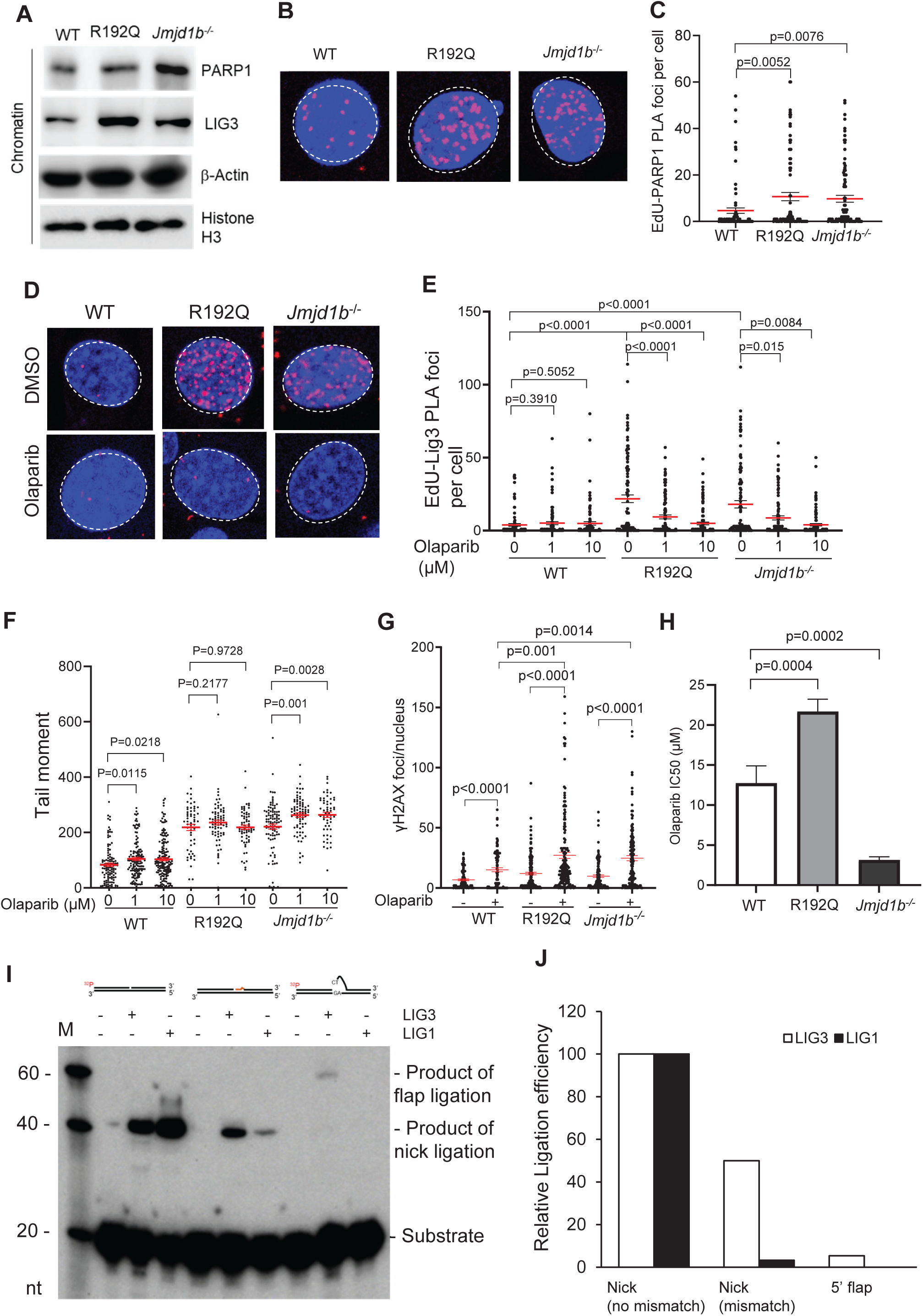
FEN1 R192Q mutation or JMJD1B deficiency induces PARP1-dependent, LIG3-mediated, error-prone OF ligation. **(A)** Chromatin associated proteins were prepared from WT, R192Q, or *Jmjd1b^-/-^*MEFs. PARP1 and LIG3 levels were determined by Western blot. β-actin and histone H3 were used as loading controls **(B, C)** PLA was performed to detect the localization of PARP1 at replication forks as represented by incorporated EdU in WT, FEN1 R192Q, and *Jmjd1b*^-/-^ MEFs. Panel B shows representative images. Panel C shows quantification of PLA puncta per cell corresponding to panel A. Data represent mean ± SEM from ≥100 cells per condition. P value from Student t-test. **(D)** PLA was performed to detect the localization of LIG3 with incorporated EdU with or without treatment with Olaparib (1 μM or 10 μM, 16h) in WT, FEN1 R192Q, and *Jmjd1b*^-/-^ MEFs. **(E)** Quantification of PLA foci per cell corresponding to panel D. Data represent mean ± SEM from ≥100 cells per condition. P value from Student t-test **. (F)** Quantification of tail moments in WT, FEN1 R192Q, and *Jmjd1b^_/_^* MEFs by alkaline BrdU comet assay (± Olaparib, 1 μM or 10 μM, 16h). Cells were pulse-labeled with 20 μM BrdU for 20 min prior to the assay. At least 50 comets were analyzed per sample. Mean tail moments ± SEMs are indicated. P value from Student t-test. **(G)** Quantification of γH2AX foci in WT, FEN1 R192Q, and *Jmjd1b^-/-^* MEFs (± Olaparib, 10 μM, 16 h) using the Image J program. (H) Viability of WT, FEN1 R192Q, and *Jmjd1b*^-/-^ MEFs treated with varying concentrations of Olaparib for 4 days was determined, IC_50_was calculated. Data represent IC_50_ values (mean ± SD, n = 4 independent assays). P value from Student t-test. **(I-J)** ^32^P-labeled DNA nick (with or without a mismatch) and 5’ flap substrates (100 nM) were incubated without or with purified LIG3 (50 nM) and LIG1 (10 nM) at 37°C for 60 min. Panel I (top) shows the diagram of different DNA substrates as specified in Supplementary Table S2 and S6 and Panel I (bottom) shows the representative 15% denaturing PAGE image. DNA substrates and products from ligation of DNA nick or 5’ flap substrates indicated. Panel J shows the quantification of the relative ligation efficiency of LIG1 or LIG3. The ligation of DNA nick substrate (no mismatch) by LIG1 (or LIG3) was set as 1. The relative ligation efficiency of other DNA substrates by LIG1 (or LIG3) was calculated accordingly.

To determine if PARP1-LIG3-mediated alternative OFM is crucial for survival of R192Q or *Jmjd1b*^-/-^ MEFs, we analyzed the viability of WT, R192Q and *Jmjd1b*^-/-^ MEFs treated with varying concentrations of Olaparib. We found that *Jmjd1b*^-/-^ MEFs were significantly more sensitive to cell killing by Olaparib than were WT MEFs (Figure 6H, Supplementary Figure S7). Surprisingly, R192Q MEFs were more resistant to PARP1 inhibitor-induced cell killing (Figure 6H, Supplementary Figure S7). To verify the distinct sensitivities of WT, R192Q, and *Jmjd1b*^-/-^ MEFs to PARP1 inhibition, we treated different MEFs with two different other selective PARP1 inhibitors BMN-673 and veliparib. Consistently, *Jmjd1b*^-/-^ MEFs were significantly more sensitive to cell killing by BMN-673 or veliparib than were WT MEFs, but R192Q MEFs were more resistant to cell killing by BMN-673 or veliparib than were WT MEFs (Supplementary Figure S8) In addition, FEN1 S187A mutation, which similarly to *Jmjd1b*^-/-^ causes persistent FEN1 association with PCNA (Guo et al., 2010), induced similar cellular phenotypes including LIG3 accumulation at chromatin (Supplementary Figure S9A), PARP1-dependent LIG3 recruitment to replication forks (Supplementary Figure S9B, S9C), and increased sensitivity to cell killing by PARP1 inhibitors (Supplementary Figure S7, S9D). This suggests that LIG1 deficiency due to persistent FEN1 association with PCNA due to defective FEN1 demethylation or phosphorylation required the PARP1-LIG3-meidated OF ligation for DNA replication and survival. However, LIG1 overexpression in R192Q MEFs may bypass the synthetic lethality from FEN1 deficiency and PARP1 inhibition as previously observed (Hanzlikova et al., 2018; He et al., 2016).

An important unanswered question is how recruitment of LIG3 to replication forks in R192Q and *Jmjd1b*^-/-^ MEFs assists in ligation of OFs that carry 5’ flap structures. To test if LIG3 could directly ligate an ssDNA flap strand to an upstream strand via microhomology DNA sequences, we prepared an oligo-based DNA nick substrate representing an OFM intermediate upon completion of FEN1-mediated 5’ flap cleavage and a flap substrate with a ssDNA flap, whose 5’ end can possibly align to the template strand via 2 microhomology sequences (Supplementary Table S2, S3). We found that LIG3, but not LIG1, could ligate the 5’ flap substrate (Figure 6I). The ligation efficiency of LIG3 on 5’ flap substrates relative to that on no-mismatch nick substrates was ∼5%, compared to 0% of LIG1 (Figure 6J). We did not detect any LIG1-mediated ligation of 5’ flap substrates, even at LIG1 concentrations as high as 15-fold (Supplementary Figure S10A). Meanwhile, we found that LIG3 but not LIG1 could effectively ligate the nick substrate bearing a DNA mismatch near the nick site (Figure 6I). The ligation efficiency of LIG3 on the mismatch-bearing nick substrates relative to that on the no-mismatch nick substrates was ∼50%, compared to 3% of LIG1 (Figure 6J). However, as the LIG1 concentration increased, the chance to ligate the DNA nick substrate with a mismatch increased remarkably (Supplementary Figure S10B). These findings suggest that recruitment of LIG3 promotes cell survival of R192Q or *Jmjd1b*^-/-^ MEFs at the expense of mutagenesis due to its activity to ligate 5’ flaps and mismatch containing DNA nicks.

### Mutations in R192Q and *Jmjd1b^-/-^* cells mapped to OF junctions define OFM-specific mutation signatures

Direct ligation of an unprocessed 5’ flap of a downstream OF to the 3’ end of an upstream OF will result in duplication, and ligation of a DNA nick with a nearby mismatch will fix a DNA point mutation. To define the frequency and spectrum of mutations in WT, R192Q, and *Jmjd1b*^-/-^ MEFs, we isolated genomic DNA and performed whole-exome sequencing (WES). We showed a considerably greater frequency of duplications, somatic single nucleotide variations (SNVs), and small insertions/deletions (Indels) in R192Q and *Jmjd1b^-/-^* MEFs than in WT MEFs (Figure 7A-7C). In addition, R192Q MEFs exhibited relatively greater duplications and SNVs than did *Jmjd1b^-/-^* MEFs, whereas *Jmjd1b^-/-^* MEFs exhibited relatively greater Indels than did R192Q (Figure 7A-7C). We compared the distributions of duplications, SNVs, and Indels with the distribution of OF junctions, which was previously defined (Li et al., 2020), in WT, R192Q, and *Jmjd1b*^-/-^ MEFs. We found that most DNA mutations in WT, R192Q, and *Jmjd1b*^-/-^ MEFs occurred near OF junctions (Figure 7D). This supports a long-standing hypothesis that OFM is a major source of DNA mutations, even in WT MEFs. In addition, we used mutation cluster analysis to reveal that SNVs in R192Q or *Jmjd1b*^-/-^ MEFs displayed Kataegis clusters, which were previously linked to DSB repair via homology-directed recombination or break-induced replication (Sakofsky et al., 2014). This is consistent with our observation that DSBs and RAD51 foci frequently occurred in R192Q or *Jmjd1b^-/-^* MEFs (Figure 5A-5D). To determine if blocking PARP1-LIG3 reduced DNA mutations in R192Q or *Jmjd1b^-/-^* MEFs, we treated them with Olaparib (5 μM, 72 h) and carried out WES. Olaparib treatment considerably reduced the frequency of duplications, but not SNVs or Indels, in R192Q or *Jmjd1b*^-/-^ compared to WT MEFs (Supplementary Figure S11).

**Figure 7.**
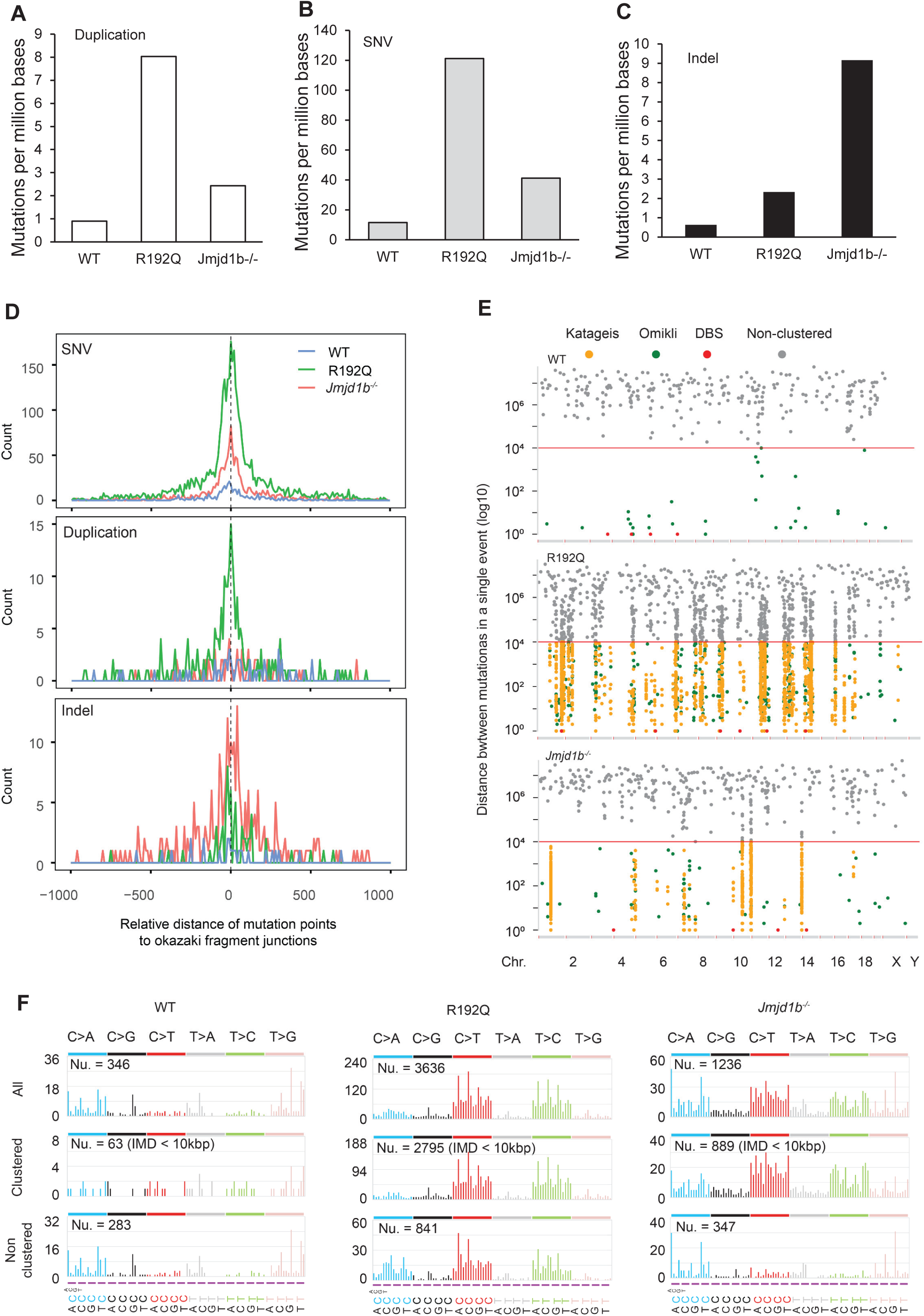
FEN1 R192Q mutation or JMJD1B deficiency leads to OFM-related duplications and SNVs. **(A-C)** WES results show the frequency of duplications (A), single nucleotide variations (SNV) (B), and small insertions/deletions (Indel (C) in WT, FEN1 R192Q, and *Jmjd1b^_/_^* MEFs. **(D)** Comparison of the distribution of duplications, SNVs, and Indel to the distribution of OF junctions in WT, FEN1 R192Q, and *Jmjd1b^_/_^* MEFs. The distance between each mutation to the nearest OF junction was calculated. **(E)** Mutation cluster analysis based on the inter-mutation distance. Kataegis (break-induced DNA synthesis), Omikli (APOBEC3-related), and DBS (doublet base substitution) clusters were defined, other events were considered non-clustered. **(F)** Mutation signatures in WT, R192Q, *Jmjd1b*^-/-^ MEFs. Occurrence of base substitutions (overall, clustered, or non-clustered) was scored based on C>A, C>G, C>T, T>A, T>C, and T>G and two flanking bases 5′ and 3′ to the mutated base.

We performed mutation signature analysis, to define whether defective FEN1 methylation (R192Q) or defective FEN1 demethylation (*Jmjd1b*^-/-^) causes a unique mutation signature and whether their mutation signatures correspond to those identified in human cancers. In R192Q MEFs, C>T or T>C mutations were considerably more frequent than other types of substitutions; in *Jmjd1b*^-/-^ MEFs, in addition to C>T or T>C, which were the most frequent mutation categories, substantial C>A mutations also occurred; in WT MEFs, C>A and T>G mutations were the most frequent (Figure 7E). In addition, we found that the mutation signatures of clustered and non-clustered mutations were similar in R192Q MEFs, but very different in the *Jmjd1b*^-/-^ MEFs (Figure 7E). These findings suggest that defects in FEN1 methylation and demethylation results in similar overall mutation signatures, partly because both induce error-prone LIG3-mediated DNA ligation.

We found that the mutation signatures in R192Q and *Jmjd1b*^-/-^ MEFs had high similarity to the clock-like SBS5 and SBS40 signature that was previously detected in most human cancers or normal cells associated with aging without a defined etiology (Supplementary Figure S12, Table S4, S5). In addition, R192Q or *Jmjd1b*^-/-^ mutation signatures were also highly similar to SBS89 that has no known etiology. It is possible that the induction of PARP1-LIG3 mutagenic OFM ligation due to R192Q or *Jmjd1b*^-/-^ are the primary cause of these mutation signatures in human cancers. R192Q or *Jmjd1b^-/-^* mutation signature was also moderately similar to the SBS44 signature previously detected in human cancers associated with mismatch repair gene deficiency (Supplementary Table S4 and S5, Supplementary Figure S12), but limited similarity to the SBS20 or SBS10a/b signatures, which were associated with Polδ or Polε mutations, respectively (Supplementary Figure S12, Table S4, S5). This suggests that the mutagenesis processes in R192Q or *Jmjd1b*^-/-^ MEFs are related to mismatch repair processes to editing out the errors from Polα but not Polδ or Polε.

## Discussion

In the current study, we define the fundamental mechanism by which PCNA coordinates the actions of Polδ, FEN1, and LIG1 for efficient and faithful OFM and DNA replication. Our studies provide evidence that Polδ, FEN1, and LIG1 sequentially rather than simultaneously bind to PCNA during OFM. In this model, Polδ and FEN1 simultaneously bind to PCNA and the Polδ-PCNA-FEN1 complex carries out displacement DNA synthesis-flap cleavage cycles, via repetitive rotary-handoff, until Polδ exits from PCNA and the OF site. The resulting PCNA-FEN1 complex catalyzes removal of the remaining 5’ flap. FEN1 then falls off PCNA, allowing recruitment of LIG1 and formation of the PCNA-LIG1 complex for ligation. Alternatively, LIG1 may be recruited to the PCNA-FEN1 complex, producing a transient FEN1-PCNA-LIG1 complex, which transforms into the PCNA-LIG1 complex for DNA ligation. Although recent biochemical and structural studies indicated that FEN1 and LIG1 can simultaneously bind to PCNA trimer subunits *in vitro* (Blair et al., 2022), our PCNA-LIG1 and FEN1-LIG1 PLA and FRET data suggest that PCNA-LIG1 is the primary functional complex for OF ligation and FEN1-PCNA-LIG1 exists only transiently. Indeed, FEN1 and LIG1 compete for binding to PCNA and DNA (Subramanian et al., 2005). We previously showed that the presence of FEN1, which cleaves DNA nicks, may result in un-ligatable DNA gaps during DNA ligation by PCNA-LIG1 (Zheng et al., 2011).

FEN1 lies at the center of this sequential PCNA binding program, and its timely association with and dissociation from PCNA are crucial for PCNA to properly recruit LIG1. FEN1 methylation dynamics are the key to these sequential processes. We previously revealed that R192 methylation mediated by PRMT5 (Guo et al., 2010) prevents FEN1 S187 phosphorylation and ensures FEN1 binding to PCNA for 5’ flap processing; conversely, FEN1 S187 phosphorylation abolishes the FEN1 interaction with PCNA. The FEN1 methylation-mimicking mutation R192F or the phosphorylation-abolishing mutation S187A result in persistent FEN1-PCNA complexes, whereas the FEN1 methylation-abolishing mutation R192K or the phosphorylation-mimicking mutation S187D cause defects in FEN1 binding to PCNA in human and mouse cells (Guo et al., 2010). Like R192K, FEN1 R192Q, a germline mutation detected in human populations, impairs FEN1 interaction with PCNA. We note that FEN1 R192Q mutation also leads to accumulation of PCNA-LIG1. This provides further evidence to support competition of FEN1 and LIG1 for PCNA binding and underscore the crucial role of FEN1 R192 methylation not only for FEN1 binding to PCNA but also for preventing premature LIG1 loading onto PCNA. Premature loading may potentially cause mutations, as it may seal the DNA nick prior to Polα error editing.

Given that FEN1 R192 methylation prevents FEN1 phosphorylation, which is essential for FEN1 dissociation from PCNA, a long-standing hypothesis has been that FEN1 is demethylated after an RNA-DNA primer is removed, allowing FEN1 to be phosphorylated by the Cdk1/Cyclin A complex (Guo et al., 2012; Guo et al., 2010; Xu et al., 2018). The fundamental question was which arginine demethylase mediates FEN1 demethylation. Our current study provides biochemical and cellular evidence demonstrating that histone arginine demethylase JMJD1B (also known as KDM3B) catalyzes FEN1 demethylation in mammalian cells. JMJD1B deficiency abolished FEN1 demethylation and dissociation from PCNA. Importantly, *Jmjd1b*^-/-^ cells, like S187A FEN1 mutant cells, showed defective LIG1 recruitment to PCNA in S phase. Intriguingly, *Jmjd1b*^-/-^ cells manage to recruit LIG1 to PCNA, resulting in a FEN1-PCNA-LIG1 complex. Because FEN1 competes with LIG1 for nick substrates, this FEN1-PCNA-LIG1 complex, unlike the PCNA-LIG complex, may potentially convert DNA nicks into un-ligatable DNA gaps. The importance of JMJD1B in facilitating OFM was further underscored by our observations that *Jmjd1b^-/-^* MEFs exhibited a high frequency of replicative SSBs, DSBs, and chromosome segregation errors.

R192Q MEFs exhibited fewer PCNA-FEN1 complexes and accumulated 5’ flaps and un-ligated OFs, likely because PCNA-FEN1 is responsible for removal of 5’ flaps and unprocessed 5’ flaps prevent OF ligation by LIG1, as shown in our biochemical assays. Intriguingly, *Jmjd1b*^-/-^ MEFs also had significantly more 5’ flaps and un-ligated OFs than the WT, although JMJD1B deficiency, opposite to R192Q, prevents FEN1 dissociation from PCNA. As persistent FEN1 interaction with PCNA delays LIG1 recruitment, OF ligation may also be delayed. Meanwhile, the failure of FEN1 to dissociate from PCNA and replication forks may lead to FEN1 competition with LIG1 for the DNA nicks between two OFs. Cleavage of the DNA nick by FEN1 resulted in DNA gaps that were not ligatable. Extension of un-ligated DNA gaps may result in 5’ flaps. A similar observation was previously reported in *Saccharomyces cerevisiae*, in which conditional depletion of DNA ligase Cdc9 resulted in accumulation of flap structures on the lagging strands of replication forks (Shi et al., 2024).

More importantly, our current studies define new mutagenic mechanisms, namely induction of error-prone PARP1-mediated OFM processes in the context of FEN1 R192Q mutation or JMJD1B deficiency. Unprocessed 5’ flaps may be converted into DSBs, the most lethal type of DNA lesion, due to unwanted nuclease cleavage of the template DNA or replication fork collapse during the next cell cycle. Therefore, to support cell survival, FEN1 R192Q or JMJD1B-deficient cells must induce alternative pathways to complete OFM and support their survival. We observed that both R192Q and *Jmjd1b*^-/-^ MEFs recruited LIG3 to replication forks. Our biochemical assays demonstrated that LIG3, unlike LIG1, can mediate ligation of 5’ ssDNA flaps to the 3’ end of the upstream DNA *in vitro*. Such enzyme activity of LIG3 is similar to that for the alternative non-homolog end joining during DSB repair (Ghezraoui et al., 2014). Our *in vitro* assays further demonstrated that LIG3 is a low-fidelity enzyme that catalyzes ligation of a DNA nick with a nearby mismatch. This error-prone nature of LIG3 contributed to increased duplications and SNVs in R192Q and *Jmjd1b*^-/-^ MEFs. It is known that proper function of the high-fidelity DNA ligase LIG1 is important to maintain accurate OFM, and LIG1 mutations could result in OFM related mutations (Williams et al., 2021) Our current study underscores the importance of that the exclusion of the low fidelity DNA ligase LIG3 at replication forks is crucial for avoidance of OFM mutation. Our studies further show that precisely regulated sequential binding of Polδ, FEN1, and LIG1 to PCNA is required for preventing LIG3 from being recruited to replication forks under normal physiology.

We demonstrated that induction of alternative LIG3-mediated OF ligation is mediated by PARP1. Inhibition of PARP1 by Olaparib abolished LIG3 recruitment to replication forks in R192Q and *Jmjd1b*^-/-^ MEFs. Interestingly, we noted a distinct impact of PARP1 inhibition on replicative SSBs and cell survival of R192Q and *Jmjd1b*^-/-^ cells: PARP1 inhibition synergized with JMJD1B deficiency in accumulating replicative SSBs and inducing cell death but did not significantly increase replicative SSBs or enhance cell death in R192Q cells. JMJD1B deficiency resulted in defective LIG1 recruitment, whereas FEN1 R192Q mutation led to the elevated expression of LIG1 and increased LIG1 interaction with PCNA. Active LIG1 in R192Q cells makes PARP1-dependent LIG3 recruitment dispensable for OF ligation and cell survival. Even though our and others’ previous studies show that FEN1 deficiency/inhibition and PARP1 inhibition synergistically kill cancer cells (Hanzlikova et al., 2018; He et al., 2016), our new observations suggest that the efficacy of combined FEN1 inhibition and PARP1 inhibition in killing cancer cells may at least partly depend on LIG1 status. Given that JMJD1B deficiency increases sensitivity to PARP1 inhibition, our results suggest a potential new mechanism by which synthetic lethality is induced for cancer treatment by coupling JMJD1B gene deficiency (e.g., 5q syndrome) or LIG1 deficiency with PARP inhibitors.

Acquisition of DNA mutations has been suggested as a major molecular mechanism that drives cancer initiation and progression. However, how such DNA mutations originate has been a long-standing question. Recent studies using next generation DNA sequencing technology on human cancer samples to address this important issue and have defined mutation signatures and further linked to many types of mutations signatures to a gene deficiency and/or exposure to certain mutagenic chemicals (Koh et al., 2021; Stratton et al., 2009). We and other groups have demonstrated that OFM, which is mediated by the sequential actions of the core enzymes Polδ, FEN1, and LIG1, is a major source of DNA mutations. However, the unique feature or signature due to defective OFM and the linkage to human cancers are undefined. In this study, we define the mutation signatures (OFM-linked signatures) in R192Q and *Jmjd1b^-/-^* MEFs, filling the important knowledge gap. Further comparison analyses reveal that the OFM-linked signatures share high similarity with the cancer mutation signatures SBS5, SBS40, and SBS89 whose aetiologic factors have not been defined. We consider that defective OFM processes may be the driving force that cause these mutation signatures in human cancers. Further studies on the mutations in OFM core enzymes or regulatory factors or induction of LIG3 mutagenic process may clearly define the etiology of these mutation signatures.

Altogether, the current work has elucidated a fundamental molecular biological mechanism of OFM as a key to mammalian DNA replication and mutation avoidance. We reveal that the arginine methylation dynamics of FEN coordinates the sequential binding of the core OFM enzymes to PCNA, which is crucial to minimizing PARP1-LIG3-induced genome-wide duplication and point mutations, which are often observed in cancer cells. Determining how the other DNA replication machinery components are post-translationally modified to meet the needs of this precise, sequential programing is warranted for future studies.

## Method details

### Formaldehyde cross-linking and preparation of soluble chromatin extraction

HeLa cells or MEFs were grown to 5 × 10^6^ cells/dish for 5 dishes. After gently washing with ice-cold PBS buffer solution, the cells were fixed with 1% formaldehyde in PBS buffer (room temperature, 10 minutes). The cross-linking reaction was quenched with addition of 0.25 M glycine/PBS in a 1:1 ratio (room temperature, 10 minutes). The fixed cells were harvested and washed twice with ice-cold PBS and were lysed in 1 ml lysis buffer (10 mM Tris-HCl, pH 7.5, 10 mM KCl, 0.5% NP-40, 0.1 mM PMSF and 1x proteinase inhibitor cocktail). After incubation on ice for 10 min, the cell suspension was centrifuged (4°C, 1, 000g, 10 min). The nuclei were suspended in a nucleus suspension buffer (10 mM Tris-HCl, pH 7.5, 150 mM NaCl, 1 mM Mg_2_Cl) containing 0.1 mM PMSF, 1x proteinase inhibitor cocktail, and 100U Benzonase). The nucleus suspension was sonicated using the Bioruptor (4°C, 10 cycles of 30s on and 30s off). The chromatin solution was incubated at 37 °C for 2 h to digest DNA and RNA molecules. NEs were kept in −80°C for storage.

### Immuno-blotting analysis of formaldehyde cross-linking chromatin extraction

Chromatin extracts were mixed with 2x SDS loading buffer in a 1:1 ratio. The samples with or without de-crosslinking by boiling for 15 minutes were resolved on an 8% SDS-PAGE gel. Proteins were transferred via a semi-dry procedure on nitrocellulose (NC) membrane, blocked for 1 h at room temperature with 5% milk powder in PBST (0.1% Tween 20 in PBS). Membranes were incubated with mouse IgG (1:2000 dilution for each IgG solution) against PCNA, FEN1, POLD1 or LIG1, respectively (4 °C, overnight). After extensively washing, the membrane was incubated HRP conjugated anti-mouse IgG (room temperature, 1 h), and detected with chemiluminescence solution (ECL, GE Healthcare).

### Proximity ligation assay (PLA)

Cells were seeded at a density of 2 × 10⁵ cells per well in 24-well plates containing sterile glass coverslips and cultured overnight in DMEM. Cells were treated with the PARP inhibitor Olaparib at the concentrations indicated in the figure legends and incubated overnight. Following treatment, cells were fixed with 4% paraformaldehyde for 30 minutes at room temperature, then permeabilized with 0.2% Triton X-100 for 15 minutes. Cells were blocked with Duolink® Blocking Solution (Sigma) for 1 h and incubated overnight at 4°C with primary antibodies diluted in Duolink® Antibody Diluent (Sigma). After washing, cells were incubated with Duolink® In Situ PLA® Probe Anti-Rabbit MINUS and Anti-Mouse PLUS (Sigma) for 1 h at 37°C. PLA reactions were performed using the Duolink® In Situ Detection Reagents Red kit (Sigma), including ligation for 30 minutes at 37°C, followed by rolling circle amplification with polymerase for 100 minutes at 37°C. PLA signals were visualized and recorded using a Zeiss Observer II or LSM 900 confocal microscope.

### Förster Resonance Energy Transfer Assay (FRET)

FRET analysis was performed to assess protein–protein interactions occurring during the OFM process. Cells were seeded at a density of 2 × 10⁵ cells per well in 24-well plates containing sterile glass coverslips and cultured overnight in DMEM. Cells were fixed, permeabilized, and blocked as described in the “Immunofluorescence” section. Following blocking, cells were incubated with primary antibodies targeting the two proteins of interest. Secondary antibodies conjugated to Alexa Fluor 568 and Alexa Fluor 647 were used to label each target protein, respectively. Alexa Fluor 568 served as the donor fluorophore, and Alexa Fluor 647 as the acceptor.

FRET imaging was performed using a Zeiss LSM 800 confocal microscope equipped with the appropriate laser lines and filter settings. Donor excitation was achieved at 561 nm, with donor emission collected at 580–600 nm. FRET signal was detected by exciting the donor (Alexa Fluor 568) at 561 nm and collecting acceptor emission (Alexa Fluor 647) at 660–700 nm. Acceptor-only images were also acquired by direct excitation at 633 nm to account for spectral bleed-through and cross-excitation. To correct for spectral overlap, control samples stained with only donor- or acceptor-labeled secondary antibodies were used to calculate bleed-through coefficients. Quantification was performed using ImageJ (NIH), applying background subtraction. Regions of interest (ROIs) were defined based on PCNA foci. The intensity of foci in the 568 nm, 647 nm, or the FRET channel that overlaps with PCNA foci were measured. The FRET intensity was corrected for bleed-through using the staining images from donor only or acceptor only (Shrestha et al., 2015).

### Liquid chromatography with tandem mass spectrometry (LC-MS/MS) analysis of protein complexes

To IP PCNA complexes from formaldehyde-fixed chromatin extracts, PCNA antibody were added to chromatin extracts in 500 μl IP buffer. The mixture was incubated at 4 °C overnight, and 50 *μl* pre-washed Pirce protein A/G magnetic beads were added and incubated room temperature for 1 h with mixing. After washing the beads with the washing buffer 3 times and with water once, the beads were incubated with a low-pH elution buffer at room temperature for 10 min to elute PCNA complexes. The PCNA complexes were mixed with 2x SDS-PAGE reducing sample buffer and the samples with or without boiling were resolved on an 8% SDS-PAGE. Protein bands were excised and proteins in complex with PCNA were analyzed using LC-MS/MS (Harvard University).

To identify FEN1-interacting proteins, HeLa cells (ATCC) stably overexpressing GFP or GFP-tagged FEN1. The absence of contamination by bacteria, yeast, and other microorganisms of the cell culture was confirmed using a mycoplasma detection kit (Sigma). The GFP or GFP-FEN1 overexpressing HeLa cells were exposed to UV radiation and NEs were prepared and GFP or GFP-tagged FEN1 and associated proteins were IPed using an anti-GFP antibody following a standard co-IP procedure as we previously described(Guo et al., 2010). The proteins in the control and FEN1 complexes were determined using LC-MS/MS (Shanghai Institute of Material Medical, Chinese Academy of Sciences, Shanghai, China).

### *In vitro* FEN1 demethylation assays

To prepare ^14^C- or ^3^H-labeled arginine methylFEN1 as the substrate for *in vitro* demethylation assays, purified recombinant 6His-tagged FEN1 (5 μg) was incubated with recombinant protein arginine symmetric dimethyl transferase PRMT5 (Yang and Bedford, 2013) (1 μg) in 30 μl of methylation buffer (50 mM HEPES (pH 8.0), 0.01% (v/v) Nonidet P-40, 10 mM NaCl, 1 mM DTT, and 1 mM PMSF) supplemented with 5 μl of S-adenosyl-L-[methyl- ^14^C]methionine or S-adenosyl-L-[methyl-^3^H]methionine (GE Healthcare) at 30°C for 2 h. 200 μl Ni-NTA agarose beads were added to the reaction. After extensive washing with PBS buffer, methylated FEN1 was eluted using imidazole (200 μM). Imidazole was removed using a Biospin 6 column (BioRad). To conduct an *in vitro* demethylation assay using purified JMJD1B, the methylated FEN1 substrate (1 μg) was incubated with purified recombinant JMJD1B (1 μg) in demethylation buffer (50 mM HEPES-KOH (pH 7.5), 1 mM 2-oxoglutarate, 2 mM ascorbate, 1 mM TCEP, 500 μM (NH_4_)_2_Fe(SO_4_)_2_·6H_2_O) in a final volume of 20 μl at 37°C for 0, 1, 2, or 4 h. The reduction of ^14^C-FEN1 or the level of released free ^3^H, which was separated from ^3^H-FEN1 using a centrifugal filter unit (cutoff 10,000 kDa), was detected, or quantitated either by autoradiography or a scintillation counter, respectively. To conduct an *in vitro* demethylation assay using HeLa NEs, the ^14^C-methylated FEN1 substrate (1 μg) was incubated with HeLa NEs with or without JMJD1B depletion (1 μg) in demethylation buffer in a final volume of 20 μl at 37 °C for 2 h. The reduction of ^14^C-FEN1 was detected by autoradiography.

### Immunoprecipitation (IP) and pull-down assays

To IP endogenous FEN1, methylated FEN1, or FEN1 complexes, mimosine-synchronized HeLa cells, primary MEFs, or bone marrow cells were collected and lysed using brief sonication in IP buffer (50 mM HEPES-KOH (pH7.4), 150 mM NaCl, 0.1% NP40, 10% glycerol, and protein inhibitor cocktail (Thermo Fisher)). After centrifugation (20,000 g, 15 min, 4°C), the supernatant was incubated with anti-FEN1 (Genetex), pan-symmetric di-methylarginine (Abcam), and protein A/G dynabeads magnetic beads (Thermo Fisher) overnight. The beads were washed three times in IP buffer, then boiled in 2x SDS-PAGE loading buffer, and resultant samples were subjected to immuno-blot analysis.

To analyze FEN1-JMJD1B physical interaction using a pull-down assay, BSA or purified recombinant FEN1 or JMJD1B was immobilized on cyanogen-bromide-activated Sepharose beads (SIGMA). 10 μl of BSA or FEN1-coated beads were incubated with 100 ng purified JMJD1B or 10 μl of BSA or JMJD1B-coated beads were incubated with 40 ng purified FEN1 in a pull-down buffer (50 mM Hepes, 100 mM NaCl, 1% bovine serum albumin, 1 mM DTT, and 10% glycerol, pH 7.5) at 4°C, overnight. After extensive washing with PBS buffer, the samples were boiled in 2x SDS-PAGE loading buffer and subjected to immuno-blot analysis.

### Immunofluorescence microscopy

The subnuclear localization sites of JMJD1B, FEN1, PCNA, LIG1, γH2AX, RAD51 were determined using indirect immunofluorescence analysis. Cells cultured on coverslips to ∼50% confluence were washed in PBS buffer and fixed in methanol at −20°C for 30 min. Immunofluorescence or co-immunofluorescence staining was carried out following a standard protocol as previously described (Guo et al., 2010). To detect JMJD1B, FEN-1, PCNA, LIG1, the fixed cells were incubated with rabbit polyclonal anti-JMJD1B (Bethyl, 1:50), mouse monoclonal anti-FEN1 (Genetex, 1:400), rabbit monoclonal anti-PCNA (Cell Signaling Technology, 1:100), mouse monoclonal anti-LIG1, rabbit monoclonal anti-γH2AX (Millipore, 1:800), or rabbit polyclonal anti-RAD51 (Abcam, 1:1000). Immunofluorescence images were analyzed and recorded using Zeiss LSM800 confocal microscope or Observer II fluorescence microscope.

### Click-iT EdU Cell Proliferation Assays

Cells were seeded at a density of 2 × 10⁵ cells per well in 24-well plates containing sterile glass coverslips and cultured overnight in the presence of the PARP inhibitor Olaparib, as indicated. To visualize subnuclear localization linked to nascent DNA synthesis, cells were incubated with 10 μM EdU for 20 minutes prior to fixation. Fixation and permeabilization were performed as described in the “Proximity Ligation Assay” section. EdU-labeled DNA was then detected by a Click reaction using a solution containing 2 mM CuSO4, 10 μM Alexa Fluor 488 azide (or azide-biotin for PLA), and 50 mM sodium ascorbate in PBS, incubated for 1 h at room temperature, protected from light when necessary. Afterward, cells were rinsed with PBS and blocked with Image-iT™ FX Signal Enhancer (Invitrogen) for 30 minutes at room temperature. Primary and secondary antibodies were diluted and applied as described in the “Immunofluorescence” section. Images were acquired using a Zeiss Observer II or LSM 900 confocal microscope.

### BrdU Comet assay

Neutral and BrdU alkaline comet assays were performed using the Comet Assay Kit (Trevigen, 4250-050). For the BrdU alkaline comet assay, cells were incubated with 20 μM BrdU. Cells were harvested and resuspended in PBS as a concentration of approximately 1X10^5^ cells/ mL. A volume of 5 μL of the cell suspension was mixed with 50 μL of 0.5% low-melting-point agarose (LMPA) at 37 °C and immediately layered onto a slide. The slides were placed at 4 °C for 10 minutes to solidify. After agarose solidification, the slides were immersed in cold lysis buffer (Trevigen, 4250-050) for at least 1 h at 4 °C. Following lysis, the slides were immersed in Alkaline Unwinding Solution (200mM NaOH, 1mM EDTA) to unwind the DNA. Electrophoresis was carried out at 21 V for 30 minutes at 4 °C in Alkaline Electrophoresis Solution (200mM NaOH, 1mM EDTA). After electrophoresis, slides were gently washed with distilled water, fixed in 70% ethanol for 10 minutes, and allowed to air dry in 37 °C. Then slides were stained with anti-BrdU (BD 347580) antibodies and secondary antibodies. Slides were imaged on a observe II microscope and ZEN 3.1 software. The tail moment of comet was measured by Open Comet of image J.

### Neutral Comet assay

Neutral and BrdU alkaline comet assays were performed using the Comet Assay Kit (Trevigen, 4250-050). The neutral comet assay was performed to evaluate DNA double-strand breaks as previously described. Briefly, cells were harvested and resuspended in PBS as a concentration of approximately 1×10^5^ cells/ mL. A volume of 5 μL of the cell suspension was mixed with 50 μL of 0.5% low-melting-point agarose (LMPA) at 37 °C and immediately layered onto a slide. The slides were placed at 4 °C for 10 minutes to solidify. After agarose solidification, the slides were immersed in cold lysis buffer (Trevigen, 4250-050) for at least 1 h at 4 °C. Following lysis, the slides were rinsed with TBE, then placed in an electrophoresis tank filled with the TBE. Electrophoresis was carried out at 21 V for 45 minutes at 4 °C. After electrophoresis, slides were gently washed with distilled water, fixed in 70% ethanol for 10 minutes, and allowed to air dry in 37 °C. The DNA was stained with SYBR Green for 30 minutes in dark. Comet images were captured using a fluorescence microscope, and tail moment were analyzed using Open Comet software in image J.

### Transmission Electron Microscopy (TEM)

Replication intermediates (RIs) were analyzed by transmission electron microscopy (TEM) following established procedures (Zellweger and Lopes, 2018) with minor modifications. Cells were treated with psoralen under 366nm UV light to crosslink and stabilize replication structures in vivo. Genomic DNA was subsequently extracted by gentle lysis, proteinase K digestion, and phenol–chloroform purification to preserve high-molecular-weight DNA with minimal shearing. Enrichment of DNA molecules containing replication structures was achieved by benzoylated-naphthoylated DEAE-cellulose chromatography. Total DNA was digested with PvuII enzyme before loading onto the column under low-salt conditions, and linear duplex DNA was removed by stepwise salt washes. Fractions enriched in single-stranded DNA–containing molecules, including replication forks and bubbles, were specifically eluted with a high-salt buffer containing caffeine. DNA was precipitated, resuspended, and subjected to buffer exchange to remove residual salts and particulates prior to grid preparation.

DNA molecules were prepared for electron microscopy using a protein-mediated spreading method adapted from classical BAC spreading procedures. Briefly, purified DNA was diluted in spreading buffer and mixed with Benzyldimethylalkylammonium chloride to promote adsorption at the air–water interface. A carbon-coated copper grid was gently touched to the surface of the droplet, allowing DNA to transfer onto the grids in an extended conformation. For contrast enhancement, the samples were rotary-shadowed with platinum at low angle in a high-vacuum evaporator. Grids were examined in a transmission electron microscope operated at 120 kV. Digital micrographs were collected at magnifications of 10,000x-20,000×. Replication intermediates were scored manually from acquired images, and structural parameters (e.g. single-stranded gaps and flaps) were quantified using image analysis software.

### In vitro DNA ligase assay

DNA ligation on the nick substrate or the 5’ flap substrate (Supplementary Table S2) was analyzed using the synthetic oligo-based DNA ligation assay system. Briefly, the upstream oligo (Upstream, Supplementary Table S3) was labeled with ^32^P at the 5’ end, and was annealed with the downstream oligo (Downstream, Downstream mismatch, or Downstream flap, Supplementary Table S3) and the template oligo (Template) to form the nick substrate (without mismatch), the nick substrate (with mismatch), and the flap substrate, respectively, following the previously published procedure (Zheng et al., 2011). Recombinant DNA ligase I (LIG1) and DNA ligase III (LIG3) proteins were incubated with the DNA substrate in buffer containing 20 mM Tris-HCl (pH 7.5), 150 mM NaCl, 10 mM MgCl2, 1 mM DTT, 50 ug/ml BSA, 0.5% Glycerol, 6 mM ATP. Reactions were carried out at 37 °C for 60 min. Products of the ligation reactions were resolved on 15% denaturing polyacrylamide gels and visualized by autoradiography.

### PARP inhibitor sensitivity assays

To determine sensitivity to Olaparib, BMN673 or Veliparib, 1 × 10^4^ WT, FEN1 R192Q, *Jmjd1b^-/-^*, or FEN1 S187A MEFs were seeded onto 12-well plates. Cells were grown in DMEM with or without varying concentrations of Olaparib, BMN673 or Veliparib (ABT-888) at 37°C for 4 days. The viable cells were counted. The cell survival rate was determined as a percentage of viable cells in each Olaparib, BMN673, or Veliparib concentration, with that of the untreated control being set as 100%.

### Whole-exome sequencing (WES) and data analysis

WES were conducted on genomic DNA isolated from WT or *Jmjd1b*^-/-^ MEFs. WES library was prepared using the KAPA DNA HyperPrep kit (Roche) with Agilent all mouse exon probes (Agilent). Libraries were sequenced on an Illumina HiSeq2500 using a paired end mode. WES sequencing reads were first assessed using FastQC (https://www.bioinformatics.babraham.ac.uk/projects/fastqc/). and filtered using Trim galore (version 0.6.10) (https://www.bioinformatics.babraham.ac.uk/projects/trim_galore/) to remove any adaptor sequence and to trim any bases with Phred quality scores lower than 20. Only paired end reads where both reads were longer than 35 bp after trimming were retained for the subsequent analysis. High-quality paired end reads from each sample were separately aligned to the reference mouse genome mm10 using Bowtie2 (version 2.4.1) (Langmead and Salzberg, 2012). The resultant sequence alignment map (SAM) file was then converted to binary alignment map (Turchi et al., 1994), sorted, and indexed using Samtools (version 1.6 using htslib 1.6). Read duplicates were removed from the sorted BAM file with MarkDuplicates from Picard toolbox (version 2.27.5) to create a BAM file with unique reads only. Single nucleotide variations or small insertions or deletions were analyzed using VarScan mpileup (v.2.4.3.1) (Koboldt et al., 2012). Germline mutations were filtered out by setting the allele frequency in the WT control as 0. The somatic mutations were scored with the mutant allele frequency in the sample no less than 0.05 and no greater than 50%. Tandem duplications and other structural variations were analyzed using Pindel (version 0.2.5b9) (Ye et al., 2009). Somatic duplications were scored if at least three supporting tracks in the sample were detected but no supporting tracks were detected in the WT control. The frequencies of the duplication or other mutations were estimated by dividing the number of somatic mutations or structure variations by size of mouse exome (∼30 millions). SigProfilerClusters v.1.0.115 (Bergstrom et al., 2022) was subsequently employed to subclassify the identified clustered mutations, incorporating a genome-wide correction for mutational density. A window size of 1 Mb was used to adjust intra-mutational distances based on local mutation density, and variant allele frequencies were also taken into account during the subclassification process. In addition, we used the SigProfilerAssignment v.0.2.56 (Díaz-Gay et al., 2023) to assign the mutation signature to reference signatures derived from Catalogue of Somatic Mutations in Cancer (COSMIC) database (Tate et al., 2019).

## Acknowledgements

We thank the Light Microscopy Digital Imaging (LMDI) Shared Resource at City of Hope for assistance with FRET analysis. We thank Sarah Wilkinson, Ph.D., for editorial assistance. Research reported in this publication included work performed in the LMDI Shared Resource supported by the National Cancer Institute of the National Institutes of Health under grant number P30 CA033572. The content is solely the responsibility of the authors and does not necessarily represent the official views of the National Institutes of Health. This work was supported by NIH grants R50 CA211397 to L.Z. and R01 CA073764 and R01 CA279840 to B.S.

## Author information

The authors declare there are no conflicts of interest in this study. Correspondence and requests for materials should be addressed to.

## Supplemental information

**Supplementary Table S1:** List of proteins in the FEN1 complexes identified by LC-MS/MS analysis.

| Rank | Protein |
| --- | --- |
| 1 | FEN1 Flap endonuclease 1 |
| 2 | PCNA Proliferating cell nuclear antigen |
| 3 | DDX46 DEAD (Asp-Glu-Ala-Asp) box polypeptide 46 (DDX46) |
| 4 | PRKDC Isoform 1 of DNA-dependent protein kinase catalytic subunit |
| 5 | POLDIP3 Polymerase delta interacting protein 46 |
| 6 | UBC;RPS27A;UBB ubiquitin and ribosomal protein S27a precursor |
| 7 | CMBL Carboxymethylenebutenolidase homolog |
| 8 | CCT2 T-complex protein 1 subunit beta |
| 9 | RPN1 Dolichyl-diphosphooligosaccharide--protein glycosyltransferase 67 kDa subunit precursor |
| 10 | VIM Vimentin |
| 11 | NAIF1 Isoform 1 of Nuclear apoptosis-inducing factor 1 |
| 12 | RNA-directed DNA polymerase (Reverse transcriptase), related domain containing protein |
| 13 | SLC1A5 Neutral amino acid transporter B(0) |
| 14 | INPP4B Type II inositol-3,4-bisphosphate 4-phosphatase |
| 15 | PRPSAP2 Phosphoribosyl pyrophosphate synthetase-associated protein 2 |
| 16 | SIPA1L2 Isoform 1 of Signal-induced proliferation-associated 1-like protein 2 |
| 17 | RBM38 14 kDa protein |
| 18 | ARSK Arylsulfatase K |
| 19 | FAM131A Isoform 2 of Protein FAM131A |
| 20 | MFN1 Isoform 2 of Mitofusin-1 |
| 21 | ADAM28 Isoform 1 of ADAM 28 |
| 22 | KCTD2 KCTD2 protein (Fragment) |
| 23 | DCD Dermcidin |
| 24 | ANXA5 Annexin A5 |
| 25 | ATP1A2 Sodium/potassium-transporting ATPase subunit alpha-2 |
| 26 | MAD1L1 Mitotic spindle assembly checkpoint protein MAD1 |
| 27 | Uncharacterized protein ENSP00000357890 (Fragment) |
| 28 | CKAP5 Isoform 1 of Cytoskeleton-associated protein 5 |
| 29 | BCAP31 B-cell receptor-associated protein 31 |
| 30 | TBR1 T-box brain protein 1 |
| 31 | SEPT1 Septin-1 |
| 32 | SRBD1 Isoform 1 of S1 RNA-binding domain-containing protein 1 |
| 33 | UACA Uveal autoantigen with coiled-coil domains and ankyrin repeats |
| 34 | MFSD9 Major facilitator superfamily domain-containing protein 9 |
| 35 | ASH1L Probable histone-lysine N-methyltransferase ASH1L |
| 36 | ZMYM5 Isoform 1 of Zinc finger MYM-type protein 5 |
| 37 | **JMJD1B Isoform 1 of JmjC domain-containing histone demethylation protein 3B** |
| 38 | FRG1 Protein FRG1 |
| 39 | GCN1L1 Translational activator GCN1 |
| 40 | EPPK1 epiplakin 1 |
| 41 | NOL6 Isoform 1 of Nucleolar protein 6 |
| 42 | MEGF8 Isoform 1 of Multiple epidermal growth factor-like domains 8 |
| 43 | GNAL guanine nucleotide binding protein (G protein), alpha activating activity polypeptide, 1 |
| 44 | LRP1 Prolow-density lipoprotein receptor-related protein 1 |
| 45 | FGD1 FYVE, RhoGEF and PH domain-containing protein 1 |
| 46 | PTPRT Isoform 1 of Receptor-type tyrosine-protein phosphatase T |
| 47 | CLTCL1 Isoform 1 of Clathrin heavy chain 2 |
| 48 | MED8 Isoform 2 of Mediator of RNA polymerase II transcription subunit 8 |
| 49 | SAMD4A SAMD4A protein |
| 50 | PRO2049 |
| 51 | NBL1 Similar to Collagen alpha-1(VIII) chain precursor |
| 52 | GP1BA platelet glycoprotein Ib alpha polypeptide precursor |
| 53 | CERK Ceramide kinase |
| 54 | RAB5B Ras-related protein Rab-5B |
| 55 | OBSCN Isoform 1 of Obscurin |
| 56 | PDIA6 Isoform 2 of Protein disulfide-isomerase A6 |
| 57 | DYNC1H1 Cytoplasmic dynein 1 heavy chain 1 |
| 58 | VPS36 Isoform 1 of Vacuolar protein-sorting-associated protein 36 |
| 59 | DLG5 Isoform 1 of Disks large homolog 5 |
| 60 | RAPGEF1 guanine nucleotide-releasing factor 2 isoform b |
| 61 | PGRMC2 cDNA FLJ34422 |
| 62 | C1orf162 Isoform 1 of Transmembrane protein C1orf162 |

\* Proteins that are also pulled down with control IgG have been excluded

**Supplementary Table 2.**
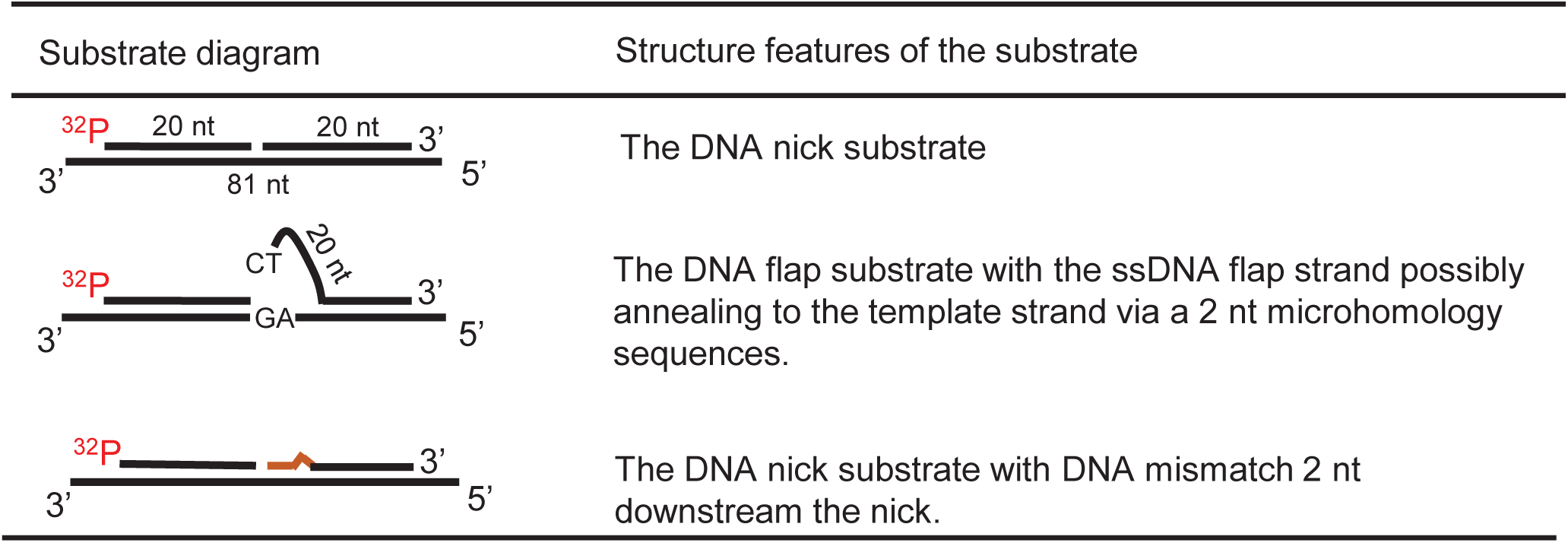
Oligo-based DNA nick or flap substrates representing different intermediates during OFM.

**Supplementary Table S3:**
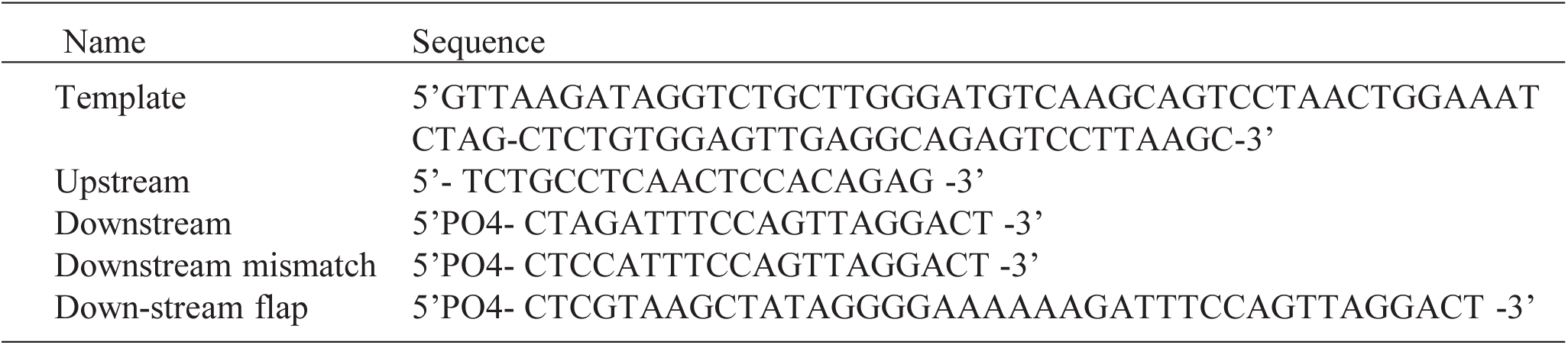
List of oligos for substrate ligation assay.

**Table S4.** The SNVs from R192Q mapping to COSMIC signatures.

| Signature | Similarity | Annotation |
| --- | --- | --- |
| SBS5 | 0.797 | clock-like signature |
| SBS40 | 0.783 | Unknown |
| SBS89 | 0.752 | Unknown |
| SBS3 | 0.734 | Tobacco smoking |
| SBS25 | 0.644 | Chemotherapy treatment |
| SBS8 | 0.563 | Unknown |
| SBS44 | 0.538 | Defective DNA mismatch repair |
| SBS18 | 0.534 | Damage by reactive oxygen species |
| SBS4 | 0.528 | Unknown (clock-like signature) |
| SBS32 | 0.51 | Azathioprine treatment |
| SBS9 | 0.502 | Polymerase eta somatic hypermutation activity |
| SBS29 | 0.493 | Tobacco chewing |
| SBS6 | 0.491 | Defective DNA mismatch repair |
| SBS39 | 0.481 | Unknown |
| SBS37 | 0.479 | Unknown |
| SBS24 | 0.471 | Aflatoxin exposure |
| SBS42 | 0.469 | Haloalkane exposure |
| SBS87 | 0.447 | Thiopurine chemotherapy treatment |
| SBS30 | 0.444 | Defective DNA base excision repair due to NTHL1 mutations |
| SBS12 | 0.44 | Unknown |
| SBS26 | 0.439 | Defective DNA mismatch repair |
| SBS36 | 0.434 | Defective DNA base excision repair due to MUTYH mutations |
| SBS31 | 0.419 | Platinum chemotherapy treatment |
| SBS35 | 0.407 | Platinum chemotherapy treatment |
| SBS15 | 0.395 | Defective DNA mismatch repair |
| SBS19 | 0.395 | Unknown |
| SBS84 | 0.392 | Activity of activation-induced cytidine deaminase (AID) |
| SBS41 | 0.38 | Unknown |
| SBS20 | 0.368 | Concurrent POLD1 mutations and defective DNA mismatch repair |
| SBS11 | 0.363 | Temozolomide treatment |
| SBS23 | 0.363 | Unknown |
| SBS1 | 0.359 | Activity of APOBEC family of cytidine deaminases |
| SBS16 | 0.349 | Unknown |
| SBS7b | 0.346 | Ultraviolet light exposure |
| SBS85 | 0.341 | Indirect effects of activation-induced cytidine deaminase (AID) |
| SBS33 | 0.317 | Unknown |
| SBS88 | 0.315 | Colibactin exposure (E.coli bacteria carrying pks pathogenicity island) |
| SBS14 | 0.311 | Concurrent polymerase epsilon mutation and defective DNA mismatch repair |
| SBS7a | 0.271 | Ultraviolet light exposure |
| SBS21 | 0.261 | Defective DNA mismatch repair |
| SBS10b | 0.25 | Polymerase epsilon exonuclease domain mutations |
| SBS86 | 0.25 | Unknown chemotherapy treatment |
| SBS7d | 0.248 | Ultraviolet light exposure |
| SBS2 | 0.218 | Defective homologous recombination DNA damage repair |
| SBS38 | 0.21 | Indirect effect of ultraviolet light |
| SBS22 | 0.206 | Aristolochic acid exposure |
| SBS7c | 0.205 | Ultraviolet light exposure |
| SBS10a | 0.199 | Polymerase epsilon exonuclease domain mutations |
| SBS28 | 0.155 | Unknown |
| SBS17a | 0.148 | Unknown |
| SBS13 | 0.128 | Activity of APOBEC family of cytidine deaminases |
| SBS34 | 0.109 | Unknown |
| SBS17b | 0.106 | Unknown |
| SBS90 | 0.09 | Duocarmycin exposure |

**Table S5.** The SNVs from *jmjd1b^-/-^* mapping to COSMIC signatures.

| Signature | Similarity | Annotation |
| --- | --- | --- |
| SBS5 | 0.805 | clock-like signature |
| SBS26 | 0.646 | Defective DNA mismatch repair |
| SBS6 | 0.63 | Defective DNA mismatch repair |
| SBS87 | 0.61 | Thiopurine chemotherapy treatment |
| SBS3 | 0.586 | Tobacco smoking |
| SBS40 | 0.58 | Unknown |
| SBS25 | 0.578 | Chemotherapy treatment |
| SBS12 | 0.575 | Unknown |
| SBS44 | 0.556 | Defective DNA mismatch repair |
| SBS1 | 0.552 | Activity of APOBEC family of cytidine deaminases |
| SBS37 | 0.499 | Unknown |
| SBS32 | 0.488 | Azathioprine treatment |
| SBS89 | 0.484 | Unknown |
| SBS15 | 0.473 | Defective DNA mismatch repair |
| SBS30 | 0.451 | Defective DNA base excision repair due to NTHL1 mutations |
| SBS31 | 0.435 | Platinum chemotherapy treatment |
| SBS42 | 0.421 | Haloalkane exposure |
| SBS9 | 0.421 | Polymerase eta somatic hypermutation activity |
| SBS33 | 0.414 | Unknown |
| SBS23 | 0.413 | Unknown |
| SBS19 | 0.409 | Unknown |
| SBS7b | 0.397 | Ultraviolet light exposure |
| SBS11 | 0.394 | Temozolomide treatment |
| SBS39 | 0.394 | Unknown |
| SBS21 | 0.393 | Defective DNA mismatch repair |
| SBS84 | 0.384 | Activity of activation-induced cytidine deaminase (AID) |
| SBS16 | 0.344 | Unknown |
| SBS20 | 0.339 | Concurrent POLD1 mutations and defective DNA mismatch repair |
| SBS24 | 0.323 | Aflatoxin exposure |
| SBS41 | 0.307 | Unknown |
| SBS7d | 0.304 | Ultraviolet light exposure |
| SBS7a | 0.302 | Ultraviolet light exposure |
| SBS35 | 0.301 | Platinum chemotherapy treatment |
| SBS4 | 0.3 | Unknown (clock-like signature) |
| SBS18 | 0.298 | Damage by reactive oxygen species |
| SBS29 | 0.298 | Tobacco chewing |
| SBS8 | 0.297 | Unknown |
| SBS17a | 0.283 | Unknown |
| SBS85 | 0.282 | Indirect effects of activation-induced cytidine deaminase (AID) |
| SBS10b | 0.262 | Polymerase epsilon exonuclease domain mutations |
| SBS88 | 0.25 | Colibactin exposure (E.coli bacteria carrying pks pathogenicity island) |
| SBS86 | 0.218 | Unknown chemotherapy treatment |
| SBS14 | 0.217 | Concurrent polymerase epsilon mutation and defective DNA mismatch repair |
| SBS36 | 0.195 | Defective DNA base excision repair due to MUTYH mutations |
| SBS2 | 0.191 | Defective homologous recombination DNA damage repair |
| SBS7c | 0.165 | Ultraviolet light exposure |
| SBS38 | 0.146 | Indirect effect of ultraviolet light |
| SBS22 | 0.127 | Aristolochic acid exposure |
| SBS10a | 0.072 | Polymerase epsilon exonuclease domain mutations |
| SBS13 | 0.057 | Activity of APOBEC family of cytidine deaminases |
| SBS34 | 0.053 | Unknown |
| SBS28 | 0.05 | Unknown |
| SBS17b | 0.049 | Unknown |
| SBS90 | 0.039 | Duocarmycin exposure |

**Supplementary Figure S1.**
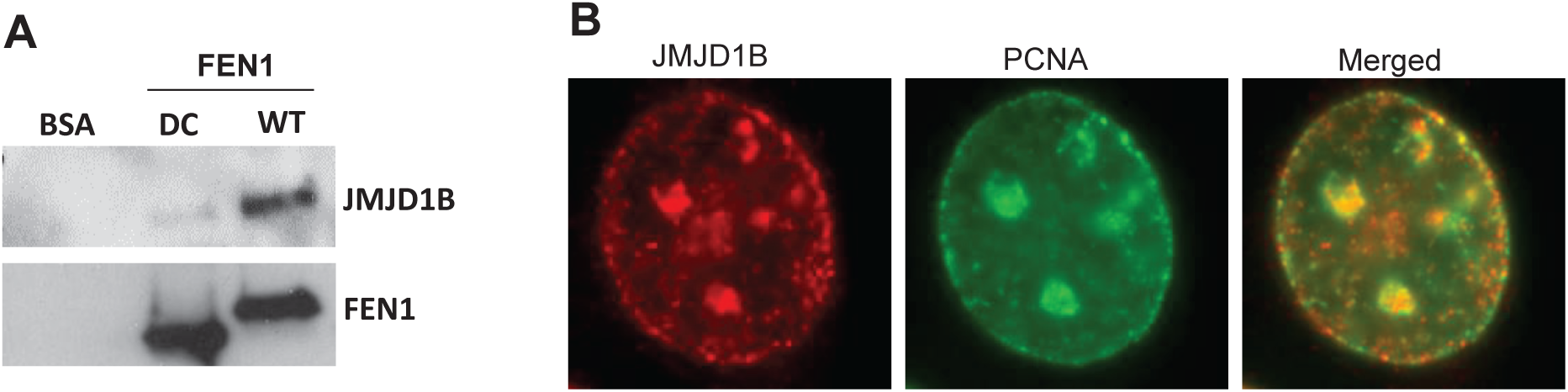
JMJD1B interacts with FEN1 and is associated with replication forks in HeLa cells. **(A)** co-pulled JMJD1B with purified WT or C-terminal deletion (DC) FEN1 proteins. **(B)** JMJD1B foci co-localizes with PCNA foci in HeLa cells.

**Supplementary Figure S2.**
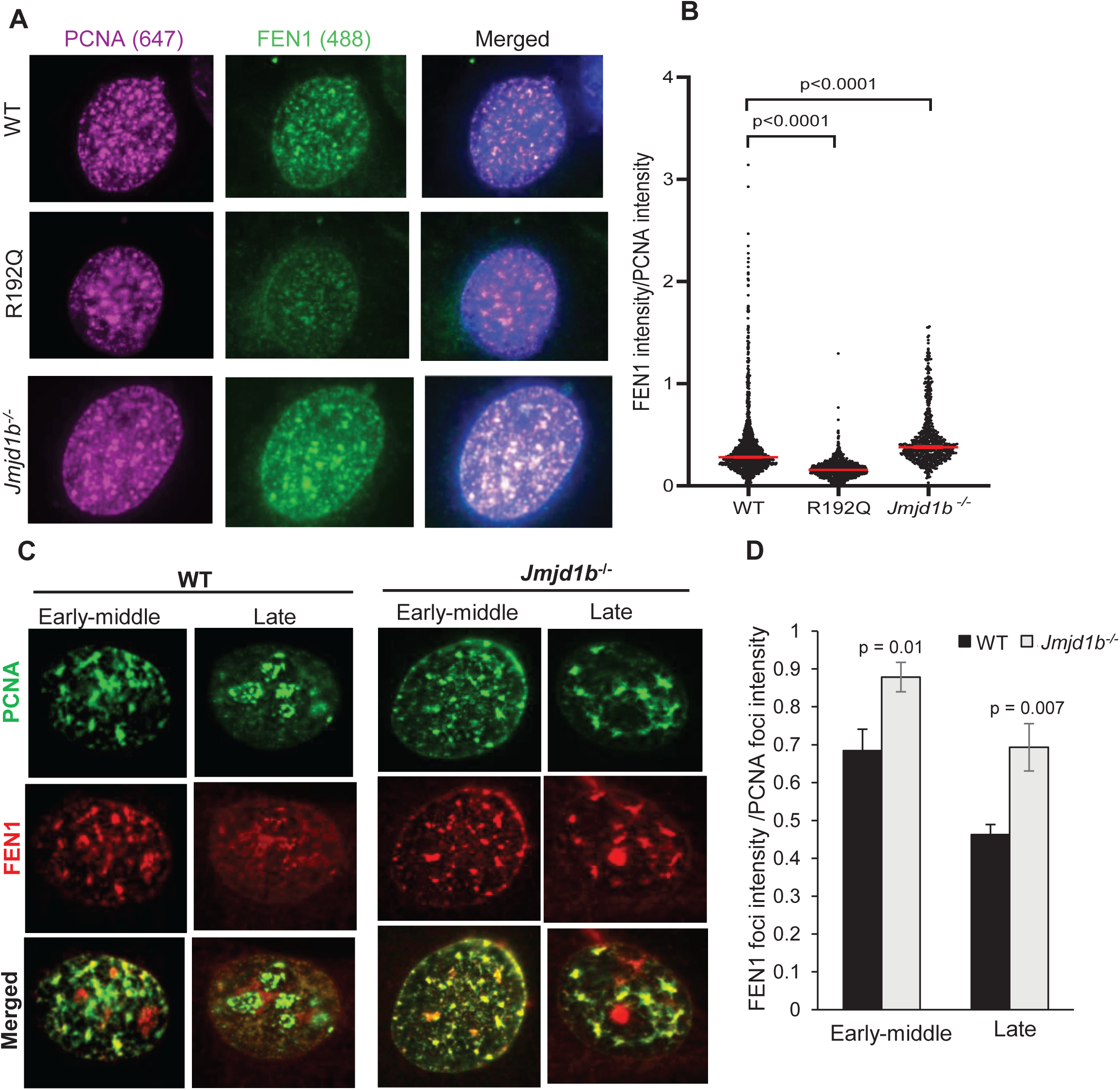
Co-IF staining shows FEN1 and PCNA co-localization in WT, R192Q, or *Jmjd1b^-/-^* MEFs. **(A)** FEN1 co-localization with PCNA in WT, R192Q and *Jmjd1b*^-/-^ MEFs. **(B)** FEN1/PCNA foci intensity ratio in WT, R192Q and *Jmjd1b*^-/-^ MEFs. Mean ± SEM are indicated. P value from Student t-test. **(C)** FEN1 co-localization with PCNA at various stages of S phase in WT and Jmjd1b-/- MEFs. **(D)** FEN1/PCNA foci intensity ratio in indicated stages of S phase in WT and Jmjd1b-/- MEFs. Mean ± SEM are indicated. P value from Student t-test.

**Supplementary Figure S3.**
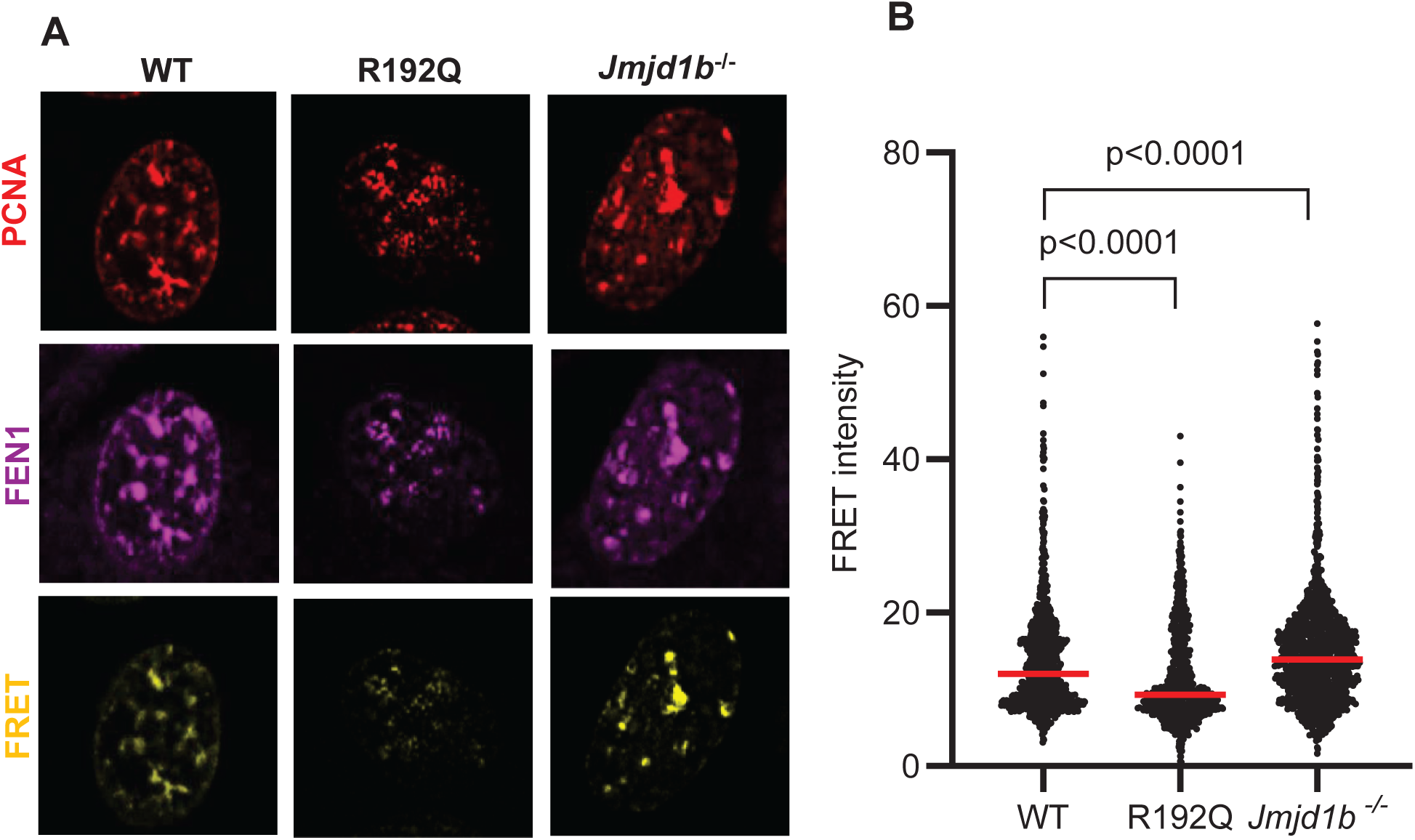
FEN1-PCNA FRET the overall level of the FEN1-PCNA complex in WT, R192Q, or *Jmjd1b^-/-^* MEFs. **(A)** representative images for the Alexa 568 (PCNA), Alexa 647 (FEN1), or the FRET channel. **(B)** The FRET intensity, which was corrected with the signals from background and bleed-through, in WT, R192Q and *Jmjd1b*^-/-^ MEFs. Mean ± SEM are indicated. P value from Student t-test.

**Supplementary Figure S4.**
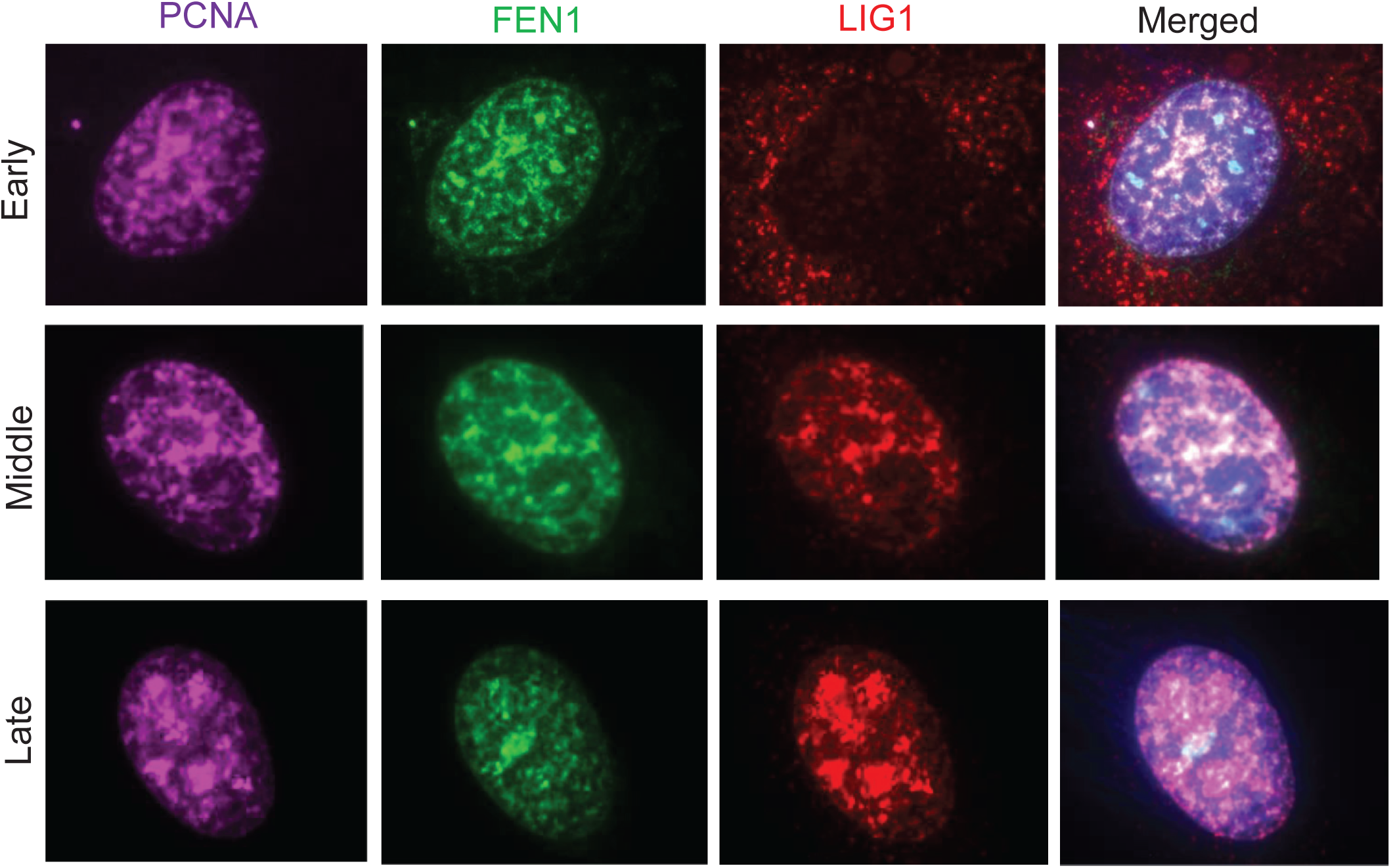
FEN1-PCNA-LIG1 co-localization in WT MEFs. Representative images for the Alexa 647 (PCNA), Alexa 488 (FEN1) and Alexa 568 (LIG1), and the merged view.

**Supplementary Figure S5.**
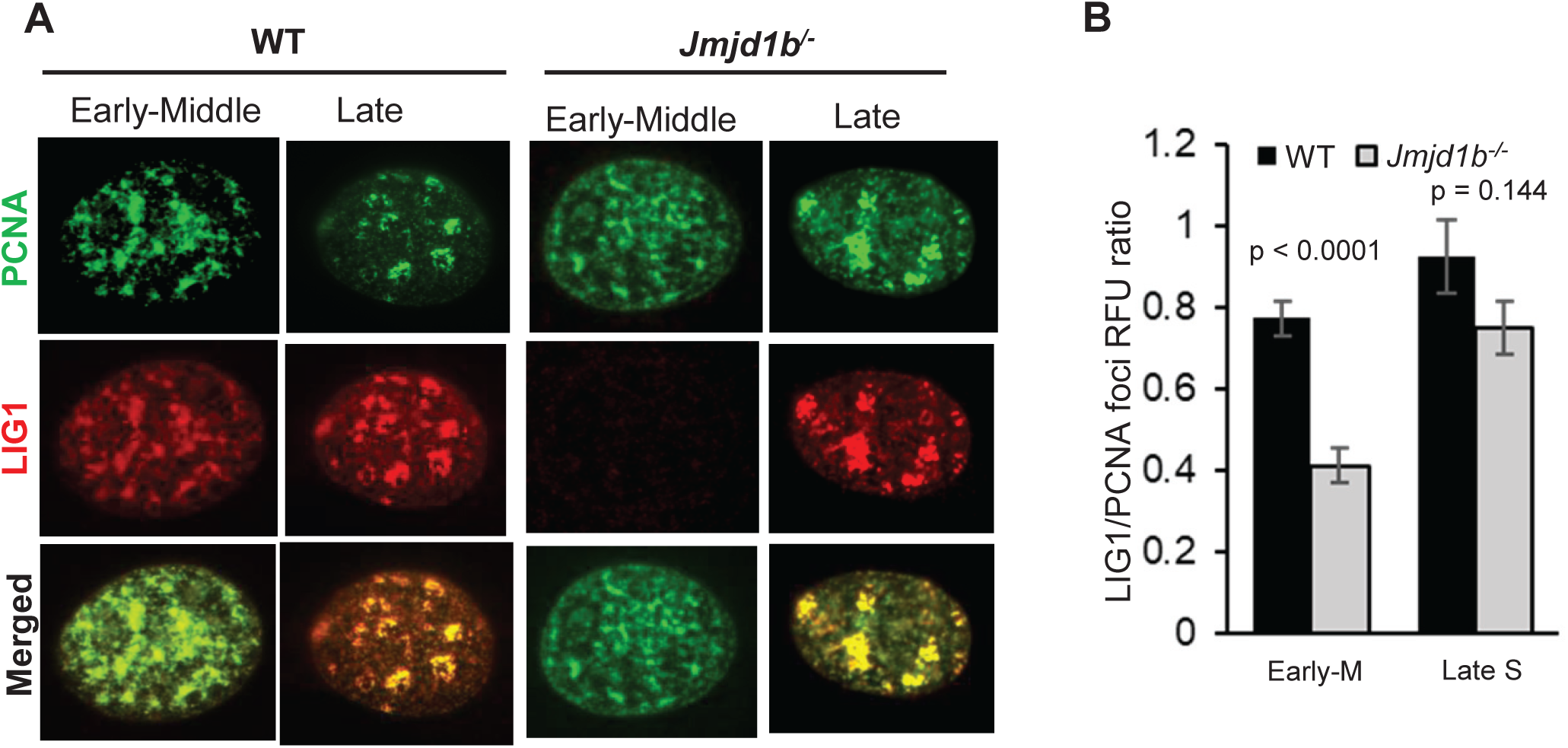
PCNA LIG1 co-localization in WT and *Jmjd1b*^-/-^ cells. **(A)** LIG1 co-localization with PCNA at various stages of S phase in WT and *Jmjd1b*^-/-^ MEFs. **(B)** LIG1/PCNA foci intensity ratio in indicated stages of S phase in WT and *Jmjd1b*^-/-^ MEFs. Mean ± SEM are indicated. P value from Student t-test.

**Supplementary Figure S6.**
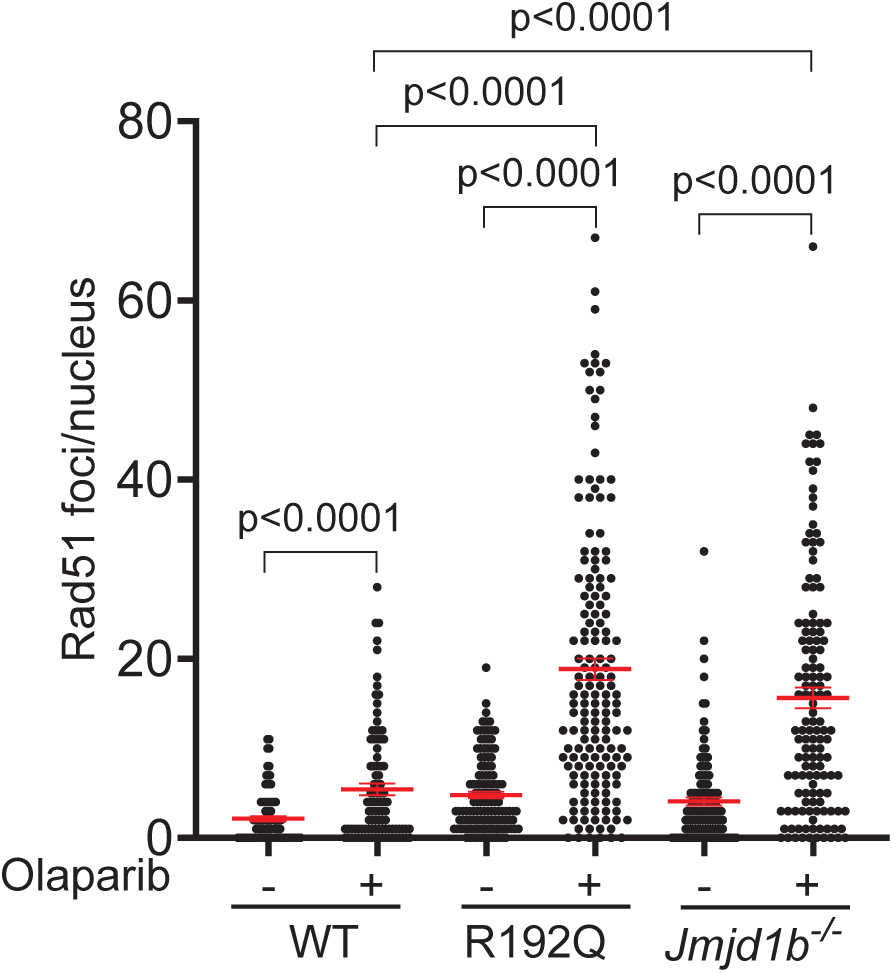
RAD51 foci in WT, R192Q, and *Jmjd1b*^-/-^ MEFs without or with Olaprib treatment. Quantification of RAD51 foci in WT, FEN1 R192Q, and *Jmjd1b^-/-^* MEFs (± Olaparib, 10 μM, 16 h) using the Image J program. Mean ± SEM. are indicated. P values are calculated using the student’s t-test.

**Supplementary Figure S7.**
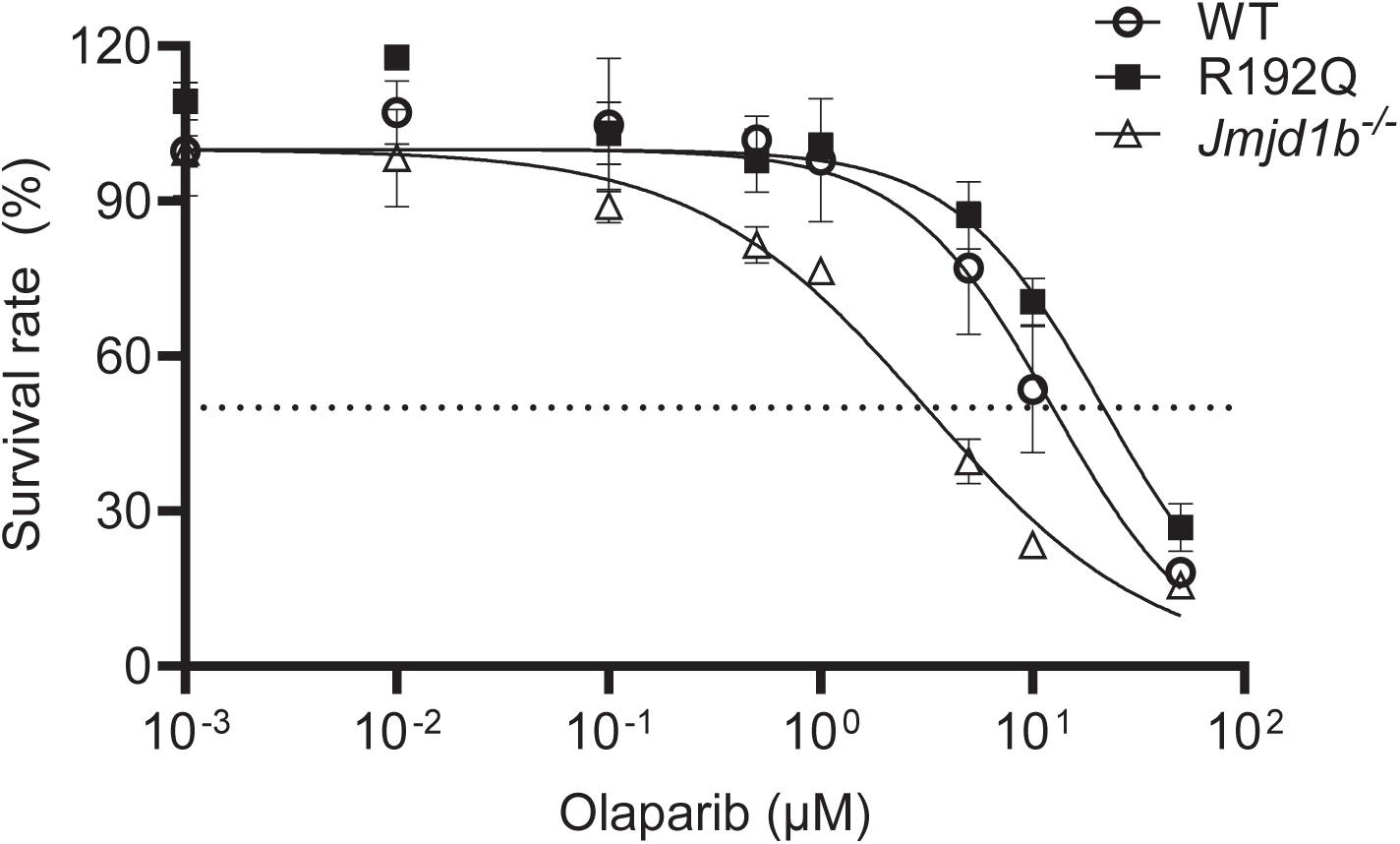
Survival curves of WT, R192Q, *Jmjd1b*^-/-^, and FEN1 S187A cells to Olaprib. Cells were seeded in 12-well plates and treated with varying concentrations of Olaparib for 96 h. Viable cells were counted and survival rate was calculated by dividing the number of surviving cells in a treatment group by those in the untreated control. Values are means ± SEM of four independent assays.

**Supplementary Figure S8.**
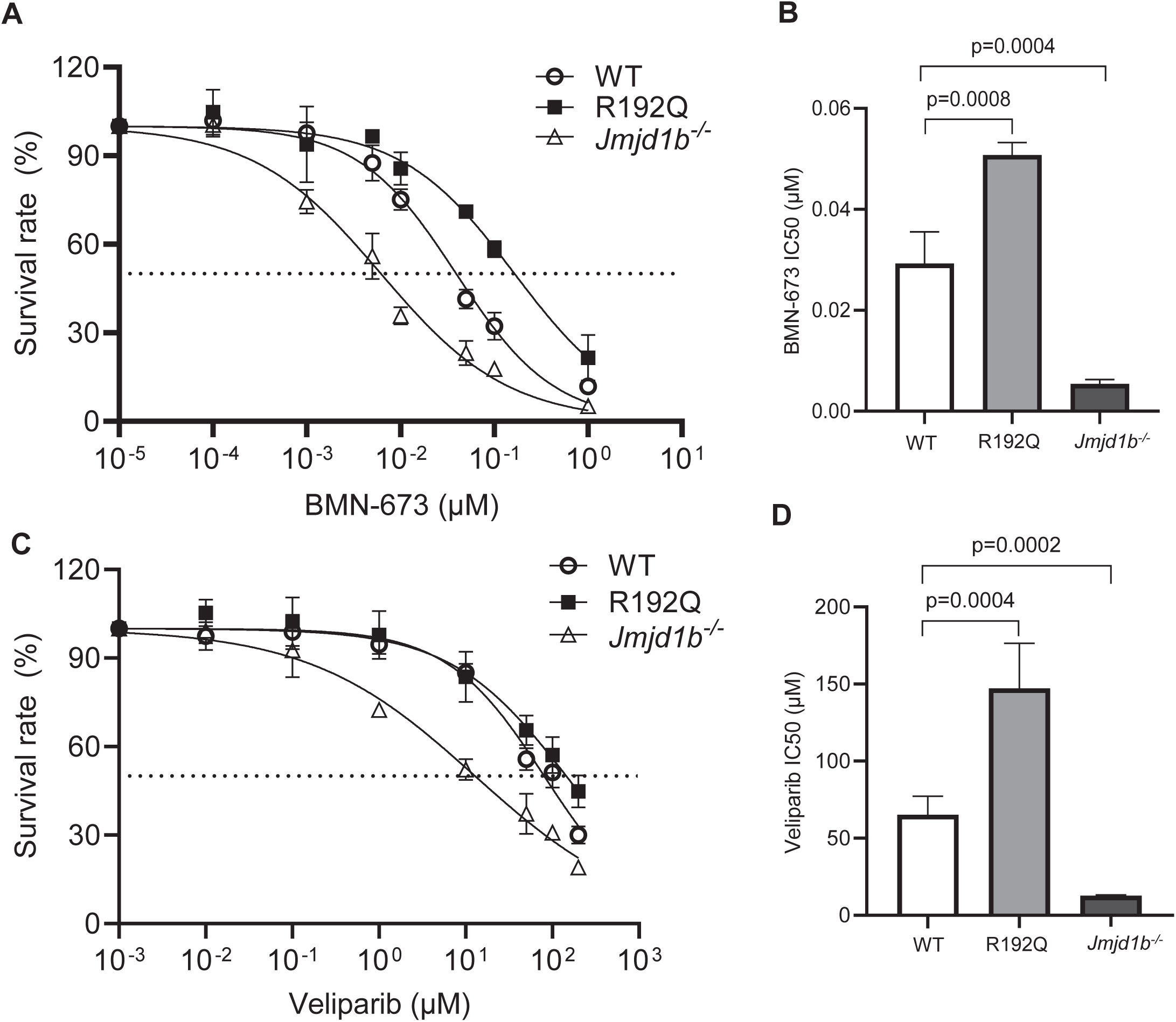
Sensitivity of WT, R192Q, *Jmjd1b*^-/-^, and FEN1 S187A cells to PARP inhibitors BMN673 or veliparib. Cells were seeded in 12-well plates and treated with varying concentrations of BMN673 (A, B), or veliparib (C,D) for 96 h. Viable cells were counted and survival rate was calculated by dividing the number of surviving cells in a treatment group by those in the untreated control. Panel A or C are survival curves of different cells treated with BMN673 or veliparib respectively. Panel B and D are the calculated IC50 based on the survival curve. Values are means ± SEM of four independent assays. P value from Student t-test.

**Supplementary Figure S9.**
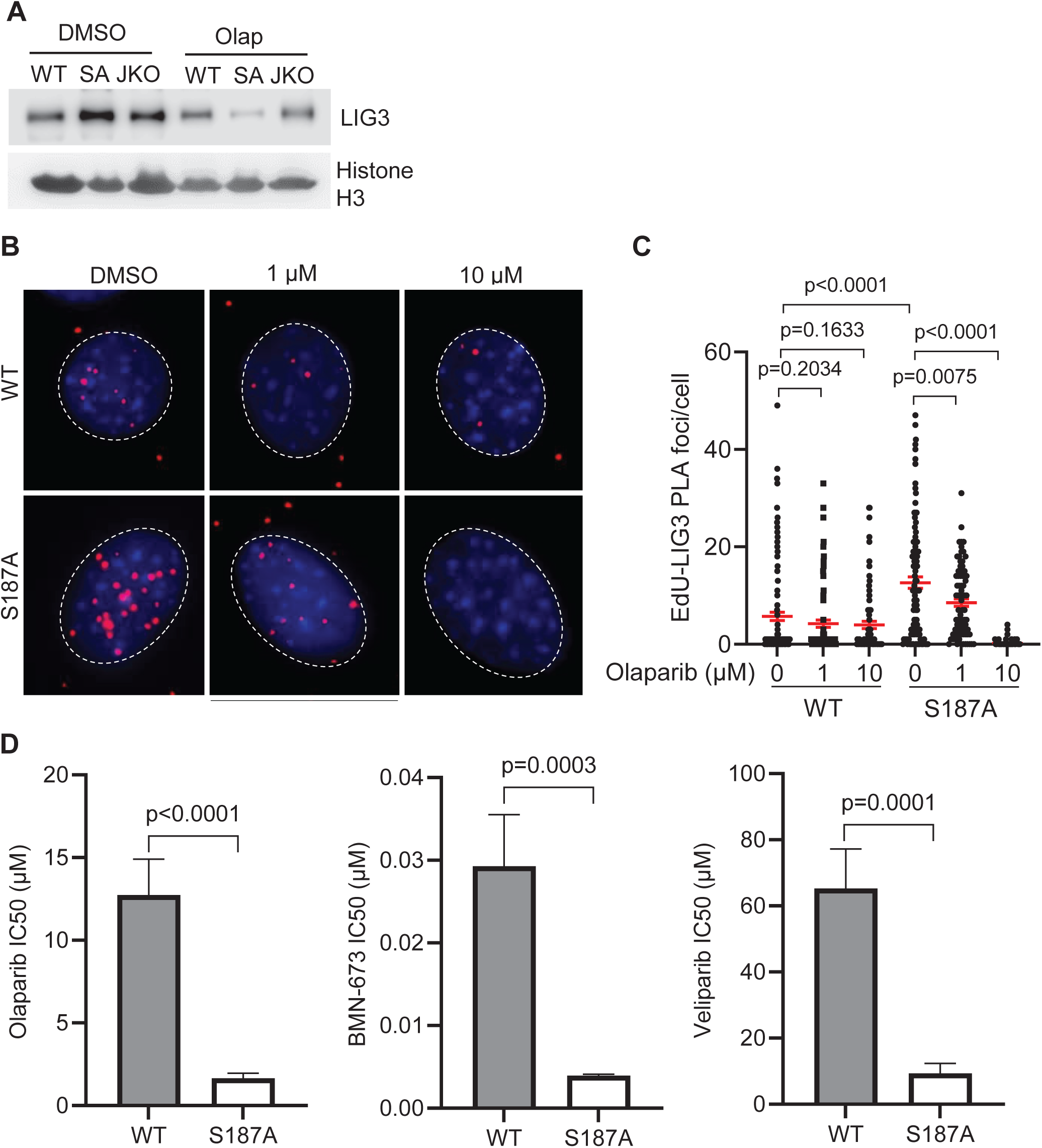
Chromatin- and replication fork-associated LIG3 in MEFs without or with Olaparib-treatment. **(A)** Western blot analysis of chromatin-associated LIG3 from WT, S187A (SA), or *Jmjd1b^-/-^* (JKO) cells treated with DMSO or Olaparib (10 μM, 16 h). Histone H3 was used as a loading control. **(B)** Representative images and **(C)** Quantification of Edu-LIG3 PLA foci in WT or S187A cells treated with DMSO or Olaparib (1, 10 μM, 16 h). Nuclei were stained with DAPI (Blue). Mean ± SEM are indicated. P value from Student t-test. **(D)** Sensitivity to cell killing of WT or S187A cells by Olaparib, BMN-673, or Veliparib. Values are mean ± SEM of four independent assays. P value from Student t-test.

**Supplementary Figure S10.**
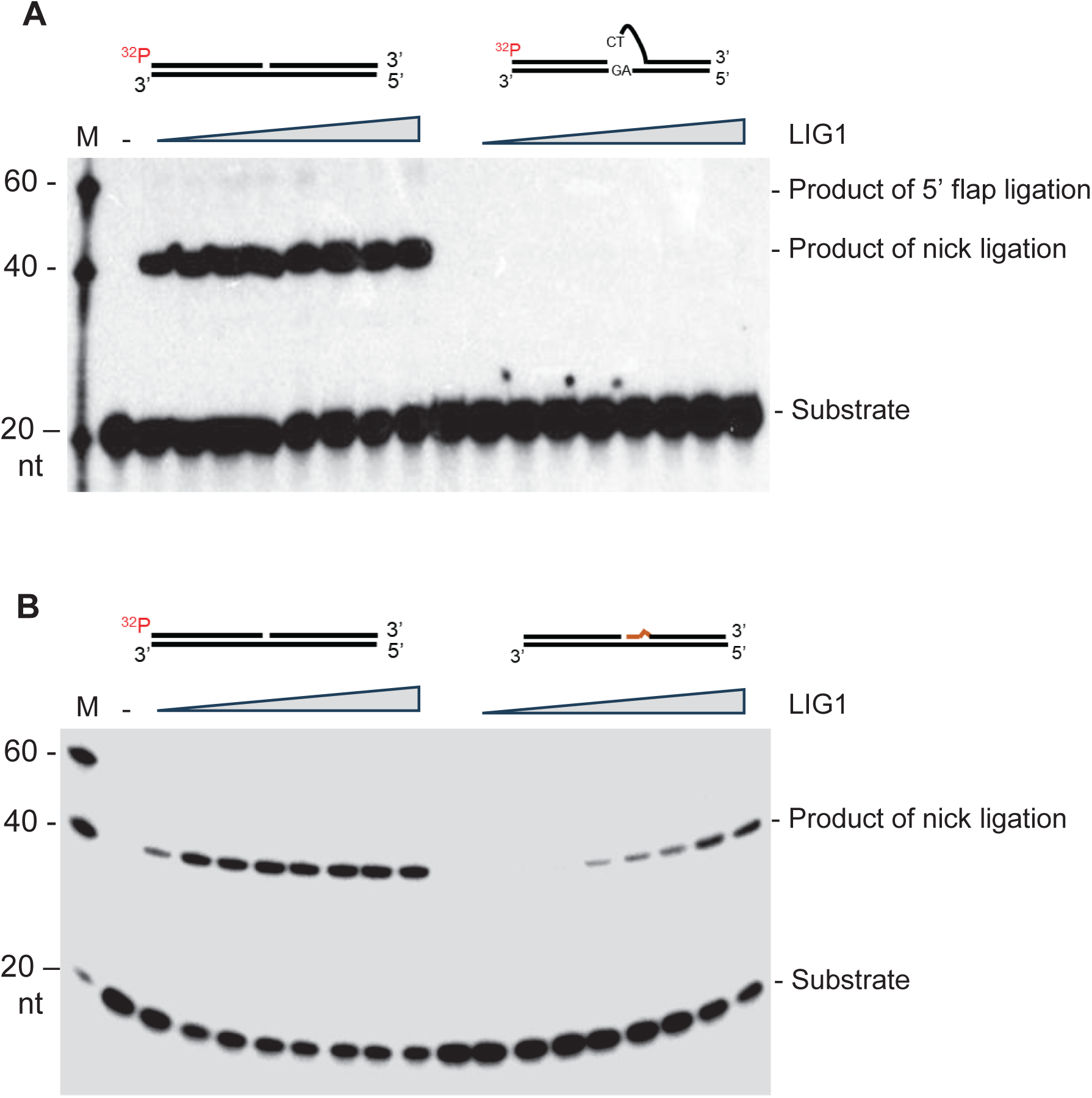
DNA ligation of DNA nick and flap substrates by LIG1. Representative denaturing PAGE image shows ligation of (A) 5’ flap substrates and (B) nick substrates by purified recombinant LIG1 of varying concentrations (5, 10, 20, 30, 40, 50, 100, 150 nM).

**Figure S11.**
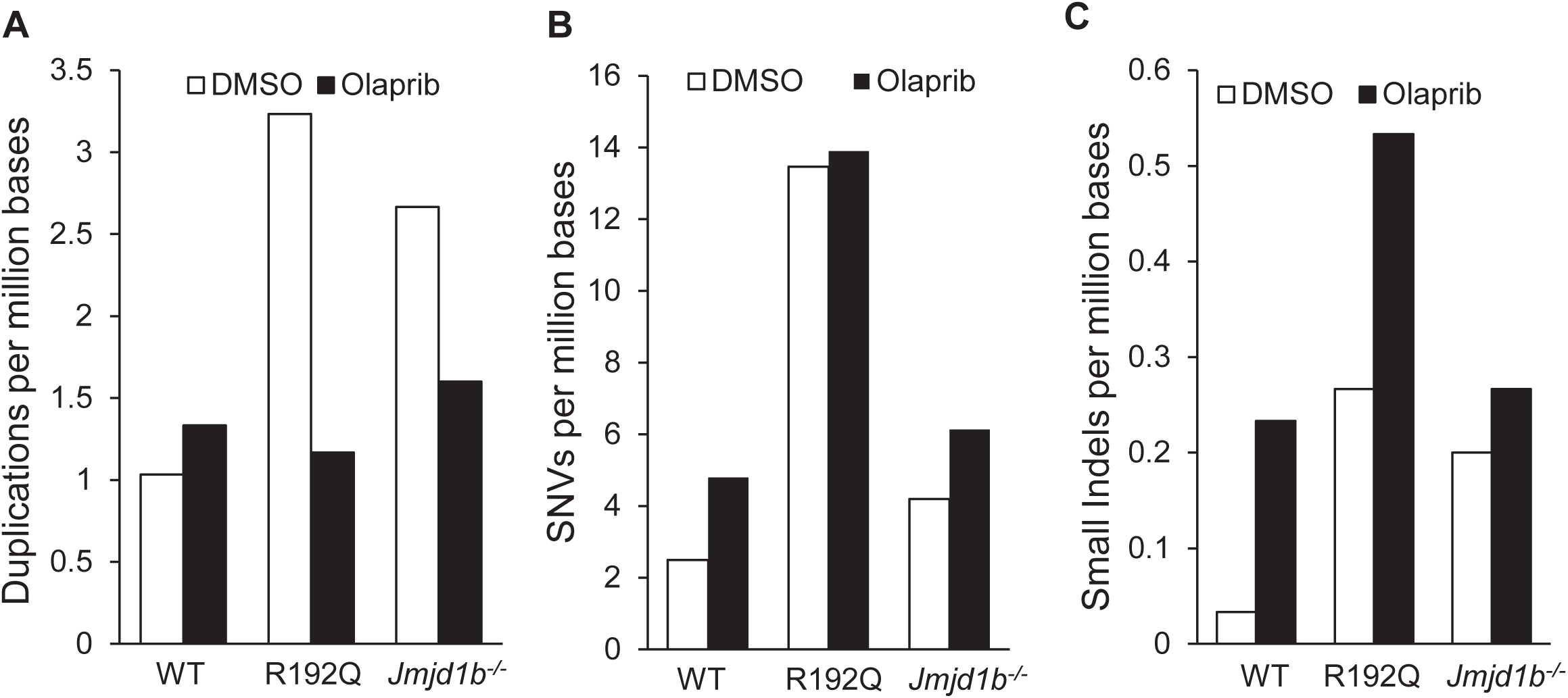
Frequency of somatic duplications, single nucleotide variation (SNV), small insertion/deletion (IN/DEL) in WT, R192Q, *Jmjd1b*^-/-^ cells with or without Olaparib treatment. Cells were treated with DMSO (control) or Olaprib for 72 hours. WES was carried out and somatic mutation frequence for (A) duplications, (B) SNVs, or (C) small Indels were calculated.

**Figure S12.**
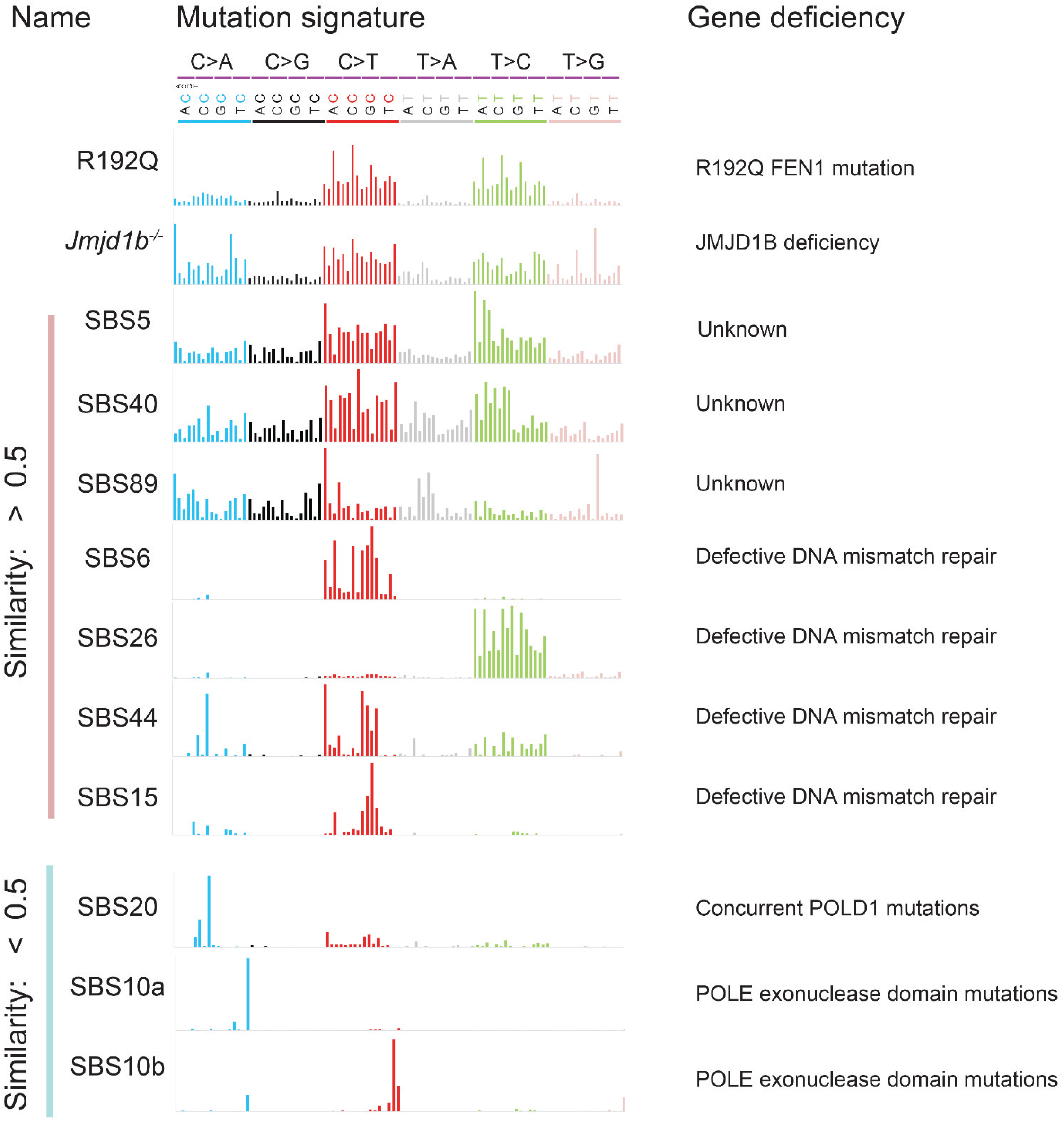
Mutation signatures in human cancers with high and low similarity to those in R192Q and *Jmjd1b^-/-^* MEFs.

## References

Abe, T., Kawasumi, R., Giannattasio, M., Dusi, S., Yoshimoto, Y., Miyata, K., Umemura, K., Hirota, K., and Branzei, D. (2018). AND-1 fork protection function prevents fork resection and is essential for proliferation. Nat. Commun 9, 3091.

Alam, M.S. (2018). Proximity ligation assay (PLA). Current protocols in immunology 123, e58.

Arakawa, H., Bednar, T., Wang, M., Paul, K., Mladenov, E., Bencsik-Theilen, A.A., and Iliakis, G. (2012). Functional redundancy between DNA ligases I and III in DNA replication in vertebrate cells. Nucleic Acids Res 40, 2599–2610.

Ayyagari, R., Gomes, X.V., Gordenin, D.A., and Burgers, P.M. (2003). Okazaki fragment maturation in yeast. I. Distribution of functions between FEN1 AND DNA2. J Biol Chem 278, 1618–1625.

Bae, S.H., Bae, K.H., Kim, J.A., and Seo, Y.S. (2001). RPA governs endonuclease switching during processing of Okazaki fragments in eukaryotes. Nature 412, 456–461.

Bergstrom, E.N., Kundu, M., Tbeileh, N., and Alexandrov, L.B. (2022). Examining clustered somatic mutations with SigProfilerClusters. Bioinformatics 38, 3470–3473.

Blair, K., Tehseen, M., Raducanu, V.-S., Shahid, T., Lancey, C., Rashid, F., Crehuet, R., Hamdan, S.M., and De Biasio, A. (2022). Mechanism of human Lig1 regulation by PCNA in Okazaki fragment sealing. Nat. Commun 13, 7833.

Budd, M.E., Reis, C.C., Smith, S., Myung, K., and Campbell, J.L. (2006). Evidence suggesting that Pif1 helicase functions in DNA replication with the Dna2 helicase/nuclease and DNA polymerase delta. Mol Cell Biol 26, 2490–2500.

Chapados, B.R., Hosfield, D.J., Han, S., Qiu, J., Yelent, B., Shen, B., and Tainer, J.A. (2004). Structural basis for FEN-1 substrate specificity and PCNA-mediated activation in DNA replication and repair. Cell 116, 39–50.

Cruz, C., Castroviejo-Bermejo, M., Gutierrez-Enriquez, S., Llop-Guevara, A., Ibrahim, Y.H., Gris-Oliver, A., Bonache, S., Morancho, B., Bruna, A., Rueda, O.M., et al. (2018). RAD51 foci as a functional biomarker of homologous recombination repair and PARP inhibitor resistance in germline BRCA-mutated breast cancer. Ann Oncol 29, 1203–1210.

Díaz-Gay, M., Vangara, R., Barnes, M., Wang, X., Islam, S.M.A., Vermes, I., Duke, S., Narasimman, N.B., Yang, T., Jiang, Z., et al. (2023). Assigning mutational signatures to individual samples and individual somatic mutations with SigProfilerAssignment. Bioinformatics 39(12).

Dieckman, L.M., Freudenthal, B.D., and Washington, M.T. (2012). PCNA structure and function: insights from structures of PCNA complexes and post-translationally modified PCNA. The eukaryotic replisome: a guide to protein structure and function, 281–299.

Dore, A.S., Kilkenny, M.L., Jones, S.A., Oliver, A.W., Roe, S.M., Bell, S.D., and Pearl, L.H. (2006). Structure of an archaeal PCNA1-PCNA2-FEN1 complex: elucidating PCNA subunit and client enzyme specificity. Nucleic Acids Res 34(*16*), 4515–4526.

Dovrat, D., Stodola, J.L., Burgers, P.M., and Aharoni, A. (2014). Sequential switching of binding partners on PCNA during in vitro Okazaki fragment maturation. Proc Natl Acad Sci U S A 111, 14118–14123.

Essers, J., Theil, A.F., Baldeyron, C., van Cappellen, W.A., Houtsmuller, A.B., Kanaar, R., and Vermeulen, W. (2005). Nuclear dynamics of PCNA in DNA replication and repair. Mol Cell Biol 25, 9350–9359.

Frank, G., Qiu, J., Somsouk, M., Weng, Y., Somsouk, L., Nolan, J.P., and Shen, B. (1998). Partial functional deficiency of E160D flap endonuclease-1 mutant in vitro and in vivo is due to defective cleavage of DNA substrates. J Biol Chem 273, 33064–33072.

Frank, G., Qiu, J., Zheng, L., and Shen, B. (2001). Stimulation of eukaryotic flap endonuclease-1 activities by proliferating cell nuclear antigen (PCNA) is independent of its in vitro interaction via a consensus PCNA binding region. J Biol Chem 276, 36295–36302.

Garg, P., Stith, C.M., Sabouri, N., Johansson, E., and Burgers, P.M. (2004). Idling by DNA polymerase delta maintains a ligatable nick during lagging-strand DNA replication. Genes Dev 18, 2764–2773.

Gary, R., Ludwig, D.L., Cornelius, H.L., MacInnes, M.A., and Park, M.S. (1997). The DNA repair endonuclease XPG binds to proliferating cell nuclear antigen (PCNA) and shares sequence elements with the PCNA-binding regions of FEN-1 and cyclin-dependent kinase inhibitor p21. J Biol Chem 272, 24522–24529.

Ghezraoui, H., Piganeau, M., Renouf, B., Renaud, J.B., Sallmyr, A., Ruis, B., Oh, S., Tomkinson, A.E., Hendrickson, E.A., Giovannangeli, C., et al. (2014). Chromosomal translocations in human cells are generated by canonical nonhomologous end-joining. Mol Cell 55, 829–842.

Guo, Z., Kanjanapangka, J., Liu, N., Liu, S., Liu, C., Wu, Z., Wang, Y., Loh, T., Kowolik, C., Jamsen, J., et al. (2012). Sequential posttranslational modifications program FEN1 degradation during cell-cycle progression. Mol Cell 47, 444–456.

Guo, Z., Zheng, L., Dai, H., Zhou, M., Xu, H., and Shen, B. (2009). Human DNA polymerase beta polymorphism, Arg137Gln, impairs its polymerase activity and interaction with PCNA and the cellular base excision repair capacity. Nucleic Acids Res 37, 3431–3441.

Guo, Z., Zheng, L., Xu, H., Dai, H., Zhou, M., Pascua, M.R., Chen, Q.M., and Shen, B. (2010). Methylation of FEN1 suppresses nearby phosphorylation and facilitates PCNA binding. Nat Chem Biol 6, 766–773.

Hanzlikova, H., Kalasova, I., Demin, A.A., Pennicott, L.E., Cihlarova, Z., and Caldecott, K.W. (2018). The importance of poly (ADP-ribose) polymerase as a sensor of unligated Okazaki fragments during DNA replication. Mol Cell 71, 319–331. e313.

He, L., Zhang, Y., Sun, H., Jiang, F., Yang, H., Wu, H., Zhou, T., Hu, S., Kathera, C.S., Wang, X., et al. (2016). Targeting DNA Flap Endonuclease 1 to Impede Breast Cancer Progression. EBioMedicine 14, 32–43.

Johnson, R.E., Kovvali, G.K., Prakash, L., and Prakash, S. (1998). Role of yeast Rth1 nuclease and its homologs in mutation avoidance, DNA repair, and DNA replication. Curr Genet 34, 21–29.

Kahli, M., Osmundson, J.S., Yeung, R., and Smith, D.J. (2019). Processing of eukaryotic Okazaki fragments by redundant nucleases can be uncoupled from ongoing DNA replication in vivo. Nucleic Acids Res 47, 1814–1822.

Koboldt, D.C., Zhang, Q., Larson, D.E., Shen, D., McLellan, M.D., Lin, L., Miller, C.A., Mardis, E.R., Ding, L., and Wilson, R.K. (2012). VarScan 2: somatic mutation and copy number alteration discovery in cancer by exome sequencing. Genome Res 22, 568–576.

Koh, G., Degasperi, A., Zou, X., Momen, S., and Nik-Zainal, S. (2021). Mutational signatures: emerging concepts, caveats and clinical applications. Nat Rev Cancer 21, 619–637.

Lancey, C., Tehseen, M., Raducanu, V.-S., Rashid, F., Merino, N., Ragan, T.J., Savva, C.G., Zaher, M.S., Shirbini, A., and Blanco, F.J. (2020). Structure of the processive human Pol δ holoenzyme. Nat. Commun 11, 1109.

Langmead, B., and Salzberg, S.L. (2012). Fast gapped-read alignment with Bowtie 2. Nat Methods 9, 357–359.

Levin, D.S., McKenna, A.E., Motycka, T.A., Matsumoto, Y., and Tomkinson, A.E. (2000). Interaction between PCNA and DNA ligase I is critical for joining of Okazaki fragments and long-patch base-excision repair. Curr Biol 10, 919–922.

Li, S., Ali, S., Duan, X., Liu, S., Du, J., Liu, C., Dai, H., Zhou, M., Zhou, L., Yang, L., et al. (2018). JMJD1B Demethylates H4R3me2s and H3K9me2 to Facilitate Gene Expression for Development of Hematopoietic Stem and Progenitor Cells. Cell Rep 23, 389–403.

Li, Z., Hua, X., Serra-Cardona, A., Xu, X., Gan, S., Zhou, H., Yang, W.S., Chen, C.L., Xu, R.M., and Zhang, Z. (2020). DNA polymerase α interacts with H3-H4 and facilitates the transfer of parental histones to lagging strands. Sci Adv 6, eabb5820.

Liu, B., Hu, J., Wang, J., and Kong, D. (2017). Direct Visualization of RNA-DNA Primer Removal from Okazaki Fragments Provides Support for Flap Cleavage and Exonucleolytic Pathways in Eukaryotic Cells. J Biol Chem 292, 4777–4788.

Ma, L., Sun, H., Abeywardana, T., Zheng, L., and Shen, B. (2022). Structure-specific nucleases: role in Okazaki fragment maturation. Trends Genet 38(*8*), 793–796.

Matsumoto, Y., Brooks, R.C., Sverzhinsky, A., Pascal, J.M., and Tomkinson, A.E. (2020). Dynamic DNA-bound PCNA complexes co-ordinate Okazaki fragment synthesis, processing and ligation. J Mol Biol 432, 166698.

Montecucco, A., Rossi, R., Levin, D.S., Gary, R., Park, M.S., Motycka, T.A., Ciarrocchi, G., Villa, A., Biamonti, G., and Tomkinson, A.E. (1998). DNA ligase I is recruited to sites of DNA replication by an interaction with proliferating cell nuclear antigen: identification of a common targeting mechanism for the assembly of replication factories. EMBO J 17, 3786–3795.

Nick McElhinny, S.A., Gordenin, D.A., Stith, C.M., Burgers, P.M., and Kunkel, T.A. (2008). Division of labor at the eukaryotic replication fork. Mol Cell 30, 137–144.

Pascal, J.M., O’Brien, P.J., Tomkinson, A.E., and Ellenberger, T. (2004). Human DNA ligase I completely encircles and partially unwinds nicked DNA. Nature 432, 473–478.

Refsland, E.W., and Livingston, D.M. (2005). Interactions among DNA ligase I, the flap endonuclease and proliferating cell nuclear antigen in the expansion and contraction of CAG repeat tracts in yeast. Genetics 171, 923–934.

Sakofsky, C.J., Roberts, S.A., Malc, E., Mieczkowski, P.A., Resnick, M.A., Gordenin, D.A., and Malkova, A. (2014). Break-induced replication is a source of mutation clusters underlying kataegis. Cell Rep 7, 1640–1648.

Sekar, R.B., and Periasamy, A. (2003). Fluorescence resonance energy transfer (FRET) microscopy imaging of live cell protein localizations. J Cell Biol 160, 629–633.

Shi, G., Yang, C., Wu, J., Lei, Y., Hu, J., Feng, J., and Li, Q. (2024). DNA polymerase delta subunit Pol32 binds histone H3-H4 and couples nucleosome assembly with Okazaki fragment processing. Sci Adv 10, eado1739.

Shrestha, D., Jenei, A., Nagy, P., Vereb, G., and Szöllősi, J. (2015). Understanding FRET as a research tool for cellular studies. Int J Mol Sci 16, 6718–6756.

Söderberg, O., Leuchowius, K.-J., Gullberg, M., Jarvius, M., Weibrecht, I., Larsson, L.-G., and Landegren, U. (2008). Characterizing proteins and their interactions in cells and tissues using the in situ proximity ligation assay. Methods 45, 227–232.

Sporbert, A., Domaing, P., Leonhardt, H., and Cardoso, M.C. (2005). PCNA acts as a stationary loading platform for transiently interacting Okazaki fragment maturation proteins. Nucleic Acids Res 33, 3521–3528.

Stith, C.M., Sterling, J., Resnick, M.A., Gordenin, D.A., and Burgers, P.M. (2008). Flexibility of eukaryotic Okazaki fragment maturation through regulated strand displacement synthesis. J Biol Chem 283, 34129–34140.

Stratton, M.R., Campbell, P.J., and Futreal, P.A. (2009). The cancer genome. Nature 458, 719–724.

Subramanian, J., Vijayakumar, S., Tomkinson, A.E., and Arnheim, N. (2005). Genetic instability induced by overexpression of DNA ligase I in budding yeast. Genetics 171, 427–441.

Tate, J.G., Bamford, S., Jubb, H.C., Sondka, Z., Beare, D.M., Bindal, N., Boutselakis, H., Cole, C.G., Creatore, C., Dawson, E., et al. (2019). COSMIC: the Catalogue Of Somatic Mutations In Cancer. Nucleic Acids Res 47, D941–d947.

Tishkoff, D.X., Filosi, N., Gaida, G.M., and Kolodner, R.D. (1997). A novel mutation avoidance mechanism dependent on S. cerevisiae RAD27 is distinct from DNA mismatch repair. Cell 88, 253–263.

Turchi, J.J., Huang, L., Murante, R.S., Kim, Y., and Bambara, R.A. (1994). Enzymatic completion of mammalian lagging-strand DNA replication. Proc Natl Acad Sci U S A 91, 9803–9807.

Waga, S., Bauer, G., and Stillman, B. (1994). Reconstitution of complete SV40 DNA replication with purified replication factors. J Biol Chem 269, 10923–10934.

Williams, J.S., Tumbale, P.P., Arana, M.E., Rana, J.A., Williams, R.S., and Kunkel, T.A. (2021). High-fidelity DNA ligation enforces accurate Okazaki fragment maturation during DNA replication. Nat Commun 12, 482.

Xu, H., Shi, R., Han, W., Cheng, J., Xu, X., Cheng, K., Wang, L., Tian, B., Zheng, L., Shen, B., et al. (2018). Structural basis of 5’ flap recognition and protein-protein interactions of human flap endonuclease 1. Nucleic Acids Res 46, 11315–11325.

Yang, Y., and Bedford, M.T. (2013). Protein arginine methyltransferases and cancer. Nat Rev Cancer 13, 37–50.

Ye, K., Schulz, M.H., Long, Q., Apweiler, R., and Ning, Z. (2009). Pindel: a pattern growth approach to detect break points of large deletions and medium sized insertions from paired-end short reads. Bioinformatics 25, 2865–2871.

Zellweger, R., and Lopes, M. (2018). Dynamic Architecture of Eukaryotic DNA Replication Forks In Vivo, Visualized by Electron Microscopy. Methods Mol Biol 1672, 261–294.

Zerjatke, T., Gak, I.A., Kirova, D., Fuhrmann, M., Daniel, K., Gonciarz, M., Muller, D., Glauche, I., and Mansfeld, J. (2017). Quantitative Cell Cycle Analysis Based on an Endogenous All-in-One Reporter for Cell Tracking and Classification. Cell Rep 19, 1953–1966.

Zheng, L., Dai, H., Hegde, M.L., Zhou, M., Guo, Z., Wu, X., Wu, J., Su, L., Zhong, X., Mitra, S., et al. (2011). Fen1 mutations that specifically disrupt its interaction with PCNA cause aneuploidy-associated cancer. Cell Res 21, 1052–1067.

Zheng, L., Dai, H., Qiu, J., Huang, Q., and Shen, B. (2007a). Disruption of the FEN-1/PCNA interaction results in DNA replication defects, pulmonary hypoplasia, pancytopenia, and newborn lethality in mice. Mol Cell Biol 27, 3176–3186.

Zheng, L., Dai, H., Zhou, M., Li, M., Singh, P., Qiu, J., Tsark, W., Huang, Q., Kernstine, K., Zhang, X., et al. (2007b). Fen1 mutations result in autoimmunity, chronic inflammation and cancers. Nat Med 13, 812–819.

Zheng, L., and Shen, B. (2011). Okazaki fragment maturation: nucleases take centre stage. J Mol Cell Biol 3, 23–30.

